# Sarbecovirus RBD indels and specific residues dictating ACE2 multi-species adaptiveness

**DOI:** 10.1101/2024.02.11.579781

**Authors:** Junyu Si, Yuanmei Chen, Mengxue Gu, Yehui Sun, Meiling Huang, Lulu Shi, Xiao Yu, Xiao Yang, Qing Xiong, Chenbao Ma, Peng Liu, Zheng-Li Shi, Huan Yan

## Abstract

Sarbecoviruses exhibit varying abilities in using angiotensin-converting enzyme 2 (ACE2) receptor^1–3^. However, a comprehensive understanding of their multi-species ACE2 adaptiveness and the underlying mechanism remains elusive, particularly for many sarbecoviruses with various receptor binding motif (RBM) insertions/deletions (indels)^4–11^. Here, we analyzed RBM sequences from 268 sarbecoviruses categorized into four RBM indel types. We extensively examined the capability of 14 representative sarbecoviruses and their derivatives in using ACE2 orthologues from 51 bats and five non-bat mammals. We revealed that most sarbecoviruses with longer RBMs (type-I), present broad ACE2 tropism, whereas viruses with single deletions in Region 1 (type-II) or Region 2 (type-III) generally exhibit narrow ACE2 tropism, typically favoring their hosts’ ACE2. Sarbecoviruses with double region deletions (type-IV) exhibit a complete loss of ACE2 usage. Subsequent investigations unveiled that both loop deletions and critical RBM residues significantly impact multi-species ACE2 tropism in different ways. Additionally, fine mapping based on type-IV sarbecoviruses elucidated the role of several clade-specific residues, both within and outside the RBM, in restricting ACE2 usage. Lastly, we hypothesized the evolution of sarbecovirus RBM indels and illustrated how loop length, disulfide, and adaptive mutations shape their multi-species ACE2 adaptiveness. This study provides profound insights into the mechanisms governing ACE2 usage and spillover risks of sarbecoviruses.

## Introduction

The Severe Acute Respiratory Syndrome (SARS) outbreak and COVID-19 pandemic significantly raised awareness of the zoonotic risks posed by sarbecoviruses^12,13^. The *Sarbecovirus* subgenus, also known as lineage B β-coronaviruses, encompasses hundreds of SARS-related viruses exhibiting varying RBM sequences^1,3–9,14–16^. Most sarbecoviruses naturally infect rhinolophus bats, the primary natural reservoir for these viruses^2,17–19^. Additionally, sarbecoviruses sharing high receptor binding domain (RBD) similarity to SARS-CoV-2 have been identified in pangolins, such as GX-P2V and GD/1/2019^9,20^. Sarbecoviruses exhibit extensive genetic diversity in RBM, likely arising from frequent recombination and the high selective pressure associated with inter-species host jumping in bats and pangolins, underscoring the risks of the emergence and outbreak of new human sarbecovirus^21–25^. However, many sarbecoviruses are known only as viral sequences and their ability to jump species and spill over to humans remains unclear.

Although ACE2 has been documented as a receptor for selected groups of setracovirus (e.g., NL63) and merbecoviruses (e.g., NeoCoV)^26,27^, it remains primarily recognized as the receptor for sarbecoviruses^1,9,24^. Notably, not all sarbecoviruses have been confirmed to use ACE2 as their receptor, especially clade 2 sarbecoviruses, which are proposed to utilize a yet unidentified receptor^1–3^. Nevertheless, ACE2 usage has been demonstrated in most representative sarbecoviruses other than clade 2 sarbecoviruses^1^. Structural analysis of ACE2 in complex with RBD from various sarbecoviruses reveals a similar interaction mode, albeit with variations in specific residues involved in recognition. The bridge-shaped RBM spanning amino acid (aa) 439-508, formed by an extended loop connecting two β strands of the RBD core subdomain and with disulfide-bridging, interacts with ACE2 through two distinct patches^28,29^. The interface on ACE2 mainly comprises the amino-terminal (N-terminal) α1-helix, along with limited interactions with the α2 helix and a loop connecting the β3 and β4 strands^28^.

Given the pivotal role of receptor recognition in governing host tropism, assessing multi-species ACE2 usage for sarbecoviruses with distinct RBM features is crucial for understanding their zoonotic potential. Previous studies have provided substantial insight into distinct receptor preferences among bats and other mammals for SARS-CoV-1, SARS-CoV-2, GX-P2V, RaTG13, NeoCoV, and others^22,24,27,30–33^. Varying entry-supporting abilities have also been observed in ACE2 orthologues from the same bat species but with different polymorphisms, particularly in residues involved in sarbecovirus binding^23,33,34^.

Sarbecoviruses are commonly classified into several clades based on the RBD phylogeny and ACE2 usage^1,3,25^. Despite sharing a similar RBD core subdomain, sarbecoviruses exhibit significant variation in RBMs, particularly the presence of various indels in Region 1 (aa443-450) or Region 2 (aa470-491) relative to SARS-CoV-2^3,35^. Clade 1 includes ACE2-using sarbecoviruses consisting of subclades 1a and 1b based on RBD phylogenetic relationships^1^. Most clade 1a (SARS-CoV-1 lineage) and 1b (SARS-CoV-2 lineage) sarbecoviruses have the longest RBM and do not carry RBM deletions^1^. Several sarbecoviruses with Region 1 or 2 single RBM deletions that are discovered phylogenetically related to clade 1b viruses, such as RshSTT182, RshSTT200^10^, Rc-o319 and Rc-kw8^4^, that recently found in Cambodia and Japan, respectively. Clade 1c, including RmYN05, RaTG15, and RsYN04, previously defined as clade 4 sarbecoviruses in some studies, were recently reported and belong to a subgroup of Asia sarbecoviruses carrying single Region 1 deletions^5,6,11^. We designated these sarbecoviruses 1c subclade considering their RBD phylogeny, geographical distribution, and ACE2 usage compared with 1a and 1b. Clade 2 sarbecoviruses are phylogenetically close to clade 1 and characterized by the presence of two deletions (indels) within the RBM^1–3,17,19,36–38^. Clade 3 sarbecoviruses, such as BM48-31, Khosta-1/2, BtKY72, and PRD-0038, discovered in Africa and Europe are considered closer to the sarbecovirus ancestors and all carry single deletions (indels) in the first RBM region (corresponding to aa443-450 of SARS-CoV-2)^1,8,35,39–41^. Several clade 3 sarbecoviruses have demonstrated ACE2 usage, suggesting it as an ancestral trait of sarbecoviruses^1,36,42,43^. Although proposed to have evolved from ACE2-using ancestors through the subsequent loss of ACE2 recognition^1,35^, whether all clade 2 sarbecoviruses have lost ACE2 usage across all ACE2 orthologues remain open.

Our understanding of the key determinants affecting sarbecoviruses ACE2 adaptiveness and the factors restricting multi-species ACE2 usage remains incomplete. With an increasing number of sarbecoviruses identified with various single RBM indels, addressing the impact of these indels on multi-species ACE2 tropism becomes crucial. Moreover, sarbecoviruses with similar RBM deletion patterns exhibit marked differences in ACE2 tropism, emphasizing the role of critical RBD residues impacting multi-species ACE2 recognition beyond loop deletions^22,23,36,44^.

In this study, we analyzed the spike sequences of 268 sarbecoviruses to delineate the overall indel features and categorized them into four RBM indel types. Employing an ACE2 library consisting of 56 orthologues, we extensively evaluated cellular RBD binding and pseudovirus entry of 14 representative sarbecoviruses and various derivatives, encompassing RBM loop chimera and mutations. Our data led to a more comprehensive understanding of the multi-species ACE2 adaptiveness across sarbecoviruses, as well as the coevolution of RBM indels and ACE2 adaptiveness.

## Results

### Four RBM indel types for sarbecoviruses

We retrieved 2318 Non-human β-coronavirus spike sequences from the NCBI and GISAID databases, with 876 distinguished as sarbecovirus based on phylogenetic analysis. After reducing redundancy by excluding identical sequences and those highly similar to SARS-CoV-1 and SARS-CoV-2 (>99% identity), we obtained 268 sarbecovirus spike sequences for subsequent investigation, consisting of 17 pangolin sarbecoviruses and 248 bat sarbecovirus, as well as 3 representative human sarbecoviruses (Fig. 1a and Supplementary data S1). Phylogenetic analysis based on RBD protein sequences revealed five sub-clades, with clade 2 accounting for the largest number (Fig. 1b, Extended Data Fig.1). Multi-sequence alignment and Sequence Logo analysis highlighted three highly variable regions in RBMs, with Regions 1 and 2, but not Region 3, being the hot spots of loop indels (Fig. 1c, Supplementary data S2).

**Fig. 1.**
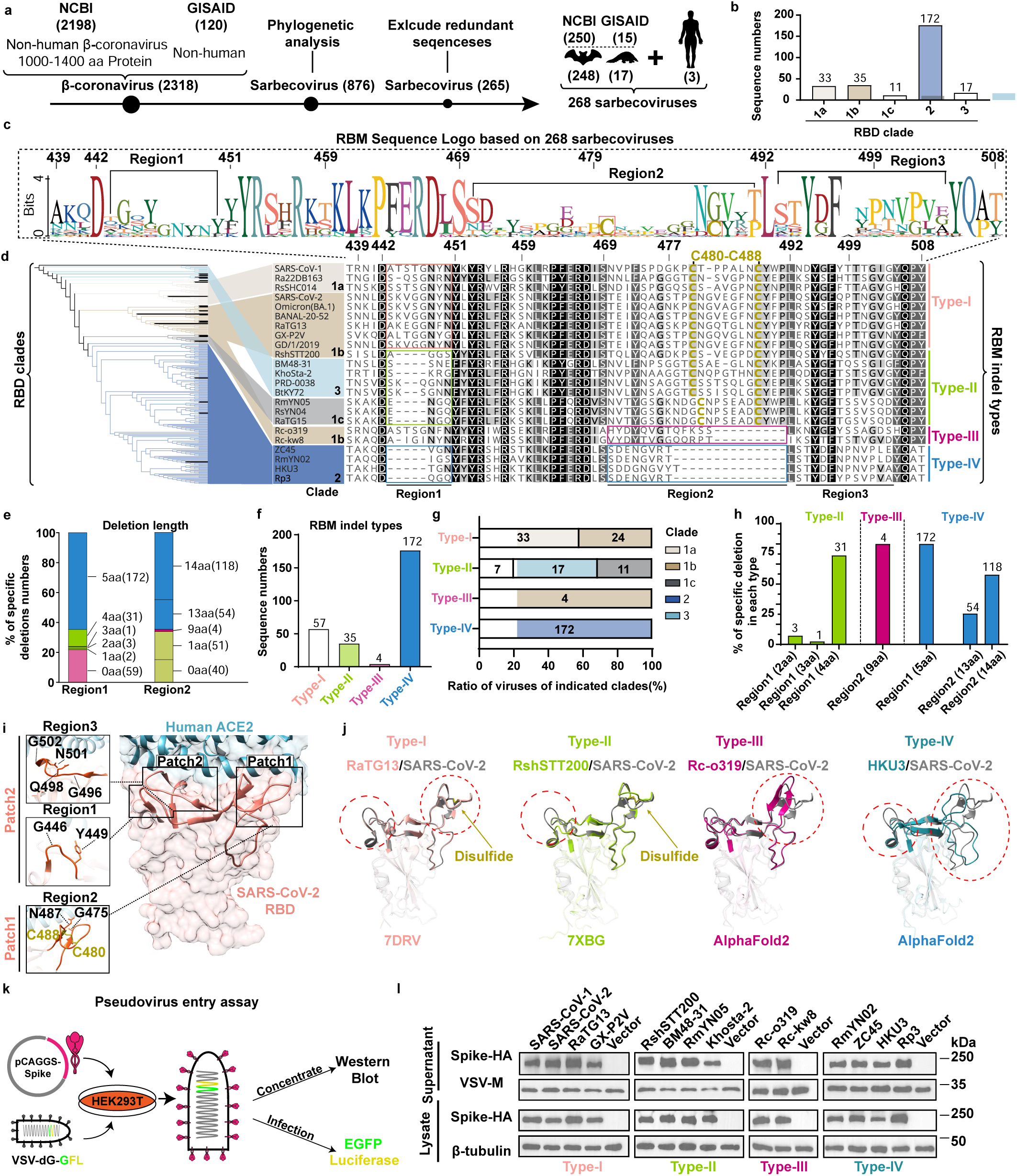
Phylogenetic and structural analysis of sarbecovirus categorized into four indel types. **a,** Flow diagram illustrating the retrieval of 265 non-redundant spike sequences from non-human sarbecovirus, and three additional human sarbecoviruses. **b,** The RBD clade information of the 268 sarbecoviruses. **c,** RBM sequence logo illustrating the three high variable regions (numbering based on SARS-CoV-2). **d,** Phylogenetic tree based on RBD amino acids for the 268 sarbecoviruses (details in Extended Data Fig. 1) and multi-sequence alignment of 23 representative sarbecoviruses with four RBM indel types. **e,** Summary of the deleted residue numbers in Region1 and Region2 compared with SARS-CoV-2 sequence. The numbers of each deletion length are indicated in the parentheses. **f,** The sequence numbers of the four RBM indel types. **g,** Distribution of RBD clades in different indel types. **h,** Analysis of the reduced length of Region1 and Region2 indels in each RBM indel types. **i,** Structural display of the two interaction patches in the SARS-CoV-2 RBD/hACE2 complex (6M0J). Residues involved in receptor recognition are indicated in the close-up views of the two interaction patches. Region 1,2 and 3 in RBD (Region 1 and 3 comprise Patch 2, while Region 2 consists of Patch 1. **j,** Structure superimposing of SARS-CoV-2 RBD (6M0J) with RBDs from representative viruses belonging to each indel types. **k,** Schematic illustration of the VSV-based pseudovirus entry assay. **l,** Western Blot detecting the level of spike protein of 14 selected sarbecoviruses in lysate or supernatant, with VSV-M and β-tubulin serving as loading controls.

RBM sequences of 23 representative sarbecoviruses spanning different clades were displayed to cover the sarbecoviruses with diversified RBM features (Fig. 1d). Amino acid identity analyses revealed that these sarbecoviruses share at least 65% spike identity and 57.84% RBD identity, whereas RBM identity can be as low as 21.54%, suggesting greater genetic variation in RBM (Extended Data Fig. 2a-c). To better investigate the impact of RBM indels on multi-species ACE2 adaptiveness, we categorized sarbecoviruses into four RBM indel types in addition to the clade-based classification. Specifically, RBM type-I describes most clade 1a and 1b sarbecoviruses with no RBM deletions (or with Region1 4aa insertions) and are considered as prototypes, RBM type-II and type-III are viruses with single RBM deletions in Region 1 or Region 2, respectively, while RBM type-IV viruses correspond to clade 2 viruses with dual RBM deletions (Fig. 1d).

Analyses of RBM deletions among the 268 sequences revealed 1 to 5 amino acid (aa) deletions in Region 1, and 1-, 9-, 13-, 14-aa deletions in Region 2 (Fig. 1e). For better classification, only deletions of 2aa or longer were applied for RBM typing. Interestingly, the 5aa deletion in Region 1 is strictly linked to 13/14aa deletions in clade 2 sarbecoviruses. This classification resulted in different sarbecovirus subgroups compared to those based on RBD clades (Fig. 1f-g). For example, all clade 1a sarbecoviruses (SARS-CoV-1 lineage) are RBM type-I, while the more complicated clade 1b (SARS-CoV-2 lineage) encompasses viruses belonging to RBM type-I, II, or III. The clade 3 and clade 1c sarbecoviruses are all grouped to RBM type-II (Fig. 1g). The length of Region 1 and 2 deletions in each RBM type is demonstrated with type-specific features (Fig. 1h).

From a structural perspective, the spatially proximate Region 1 and Region 3 loops form interaction patch 2, while the majority of residues in Region 2 loop contribute to interaction patch 1 (Fig. 1i). Interestingly, cysteine residues are rare in RBM, with only one highly conserved disulfide bridge for stabilizing loop in Region 2, which is absent in RBM type III and IV sarbecoviruses (Fig. 1d, i)^45^. Superimposition of the solved or AlphaFold2-predicted RBDs with that of SARS-CoV-2 highlighted notable differences in the extended loops carrying specific deletions (Fig. 1j). Given that the two deletions are situated in critical RBM extensions for ACE2 interaction, their presence is considered to impact multi-species ACE2 adaptiveness.

Given the unavailability of the authentic sarbecovirus strains, we employed a dual reporter-based vesicular stomatitis virus (VSV) pseudotyping system carrying sarbecovirus spikes to assess receptor functionality of various ACE2 orthologues (Fig. 1k, Extended Data Fig. 3a-c)^3^. The spike proteins from these sarbecoviruses were successfully incorporated into the VSV pseudotypes at comparable levels (Fig. 1l). In addition, a well-established RBD-hFc-based assay was also employed to assess the live cell virus-receptor binding (Extended Data Fig. 3d-f). The two different functional assays provide cross-validation and, to an extent, exclude the potential impact of other spike components on viral entry, such as NTD and S2.

### Multi-species ACE2 usage profile

To illustrate a comprehensive ACE2 usage spectrum of each sarbecovirus, we examined 56 ACE2 orthologues from 51 bats and 5 representative non-bat mammals. The bat species represent a broad genetic diversity spanning 11 bat families with global distribution, including eight rhinolophus bats geographically across Europe, Africa, and Asia (Fig 2a, and Extended Data Fig. 4)^30^. Sequence analysis of these ACE2 orthologues exhibited great diversity in residues potentially involved in sarbecovirus interactions (Extended Data Fig.5a-b). HEK293T cells stably expressing ACE2 orthologues were established and maintained with verified expression^30^ (Extended Data Fig. 6). RBD binding and pseudovirus entry assays were conducted to evaluate the multi-species ACE2 usage of 14 sarbecoviruses with distinct RBM features (Fig. 2a-b).

**Fig. 2.**
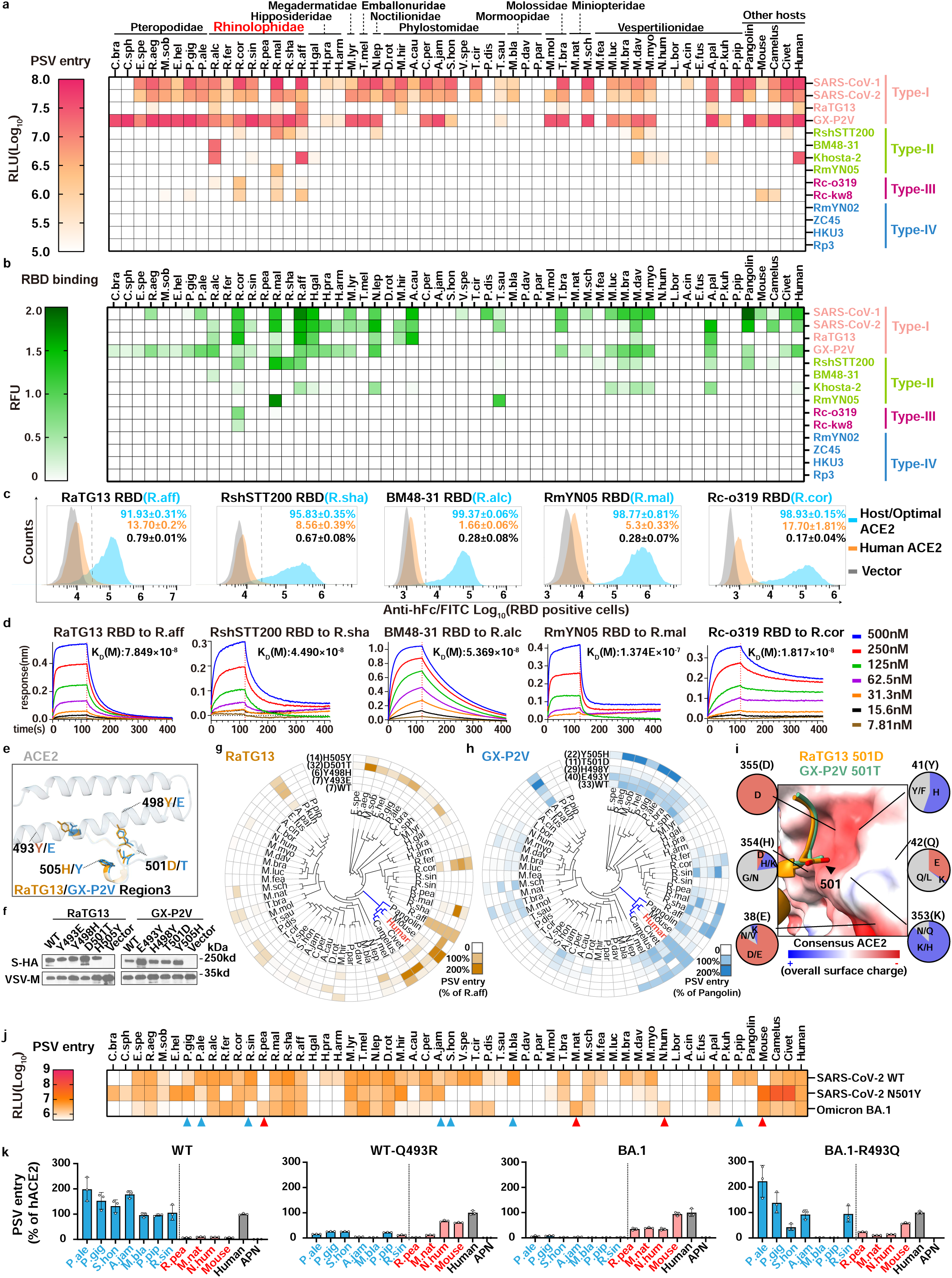
Multi-species ACE2 tropism of 14 representative sarbecoviruses and the contribution of critical RBM residues. **a, b**, Multi-species ACE2 usage spectra of sarbecoviruses of different indel types. PSV entry **(a)** and RBD binding **(b)** of 14 sarbecoviruses based on HEK293T cells stably expressing the 56 ACE2 orthologues from bats and selected mammals **c**, **d**, RBD binding efficiencies of selected sarbecoviruses favoring their hosts’ ACE2(or the optimal ACE2) analyzed by Flow cytometry (**c**) or Biolayer interferometry (BLI) (**d**). **e**, **f**, Structural demonstration (PDB IDs: 7DRV, 7DDP) (**e**) and spike protein packaging efficiencies (**f**) of RaTG13 and GX-P2V swap mutants. **g, h,** Heat map displaying PSV entry efficiencies of RaTG13 and GX-P2V swap mutants in HEK293T cells expressing the indicated ACE2 orthologues. PSV entry > 5% is considered as an effective entry and the number of ACE2 support entry is showed in parentheses (SARS-CoV-2 numbering). **i,** Negative charged surface of the consensus ACE2 (based on 56 ACE2 orthologues) spatially proximate to residue 501 of RaTG13 and GX-P2V. The structure of consensus ACE2 is predicted by AlphaFold2, and the interaction is predicted by HDOCK. **j,** PSV entry efficiencies of SARS-CoV-2 WT, N501Y, and Omicron BA.1 in HEK293T stably expressing the 56 ACE2s orthologues. Red triangle: increased efficiencies to support Omicron; Blue triangle: reduced efficiency to support Omicron. **k**, PSV entry efficiencies of SARS-CoV-2 mutants in HEK293T stably expressing the indicated ACE2 orthologues. RLU: Relative Luminescence Units; RFU: Relative Fluorescence Units. The amino acid usage of the residues consisting of the surface are indicated. Data representative of 2-3 independent experiments for a, b, g, h, j, and k (n=3 biological replicates). Mean ± SD for k.

These two assays displayed generally consistent ACE2 usage patterns, with a few exceptions. Except for the type-IV RBM sarbecoviruses, the other ten sarbecoviruses displayed ACE2 usage with different tropisms. Type-I RBM viruses, like SARS-CoV-1, SARS-CoV-2, and GX-P2V, efficiently use most orthologues, including human ACE2 (hACE2) (Fig.2a). Comparatively, RBM type-II or type-III sarbecoviruses generally showed narrower ACE2 tropism, and most are unable to use hACE2. The geographical distributions of rhinolophus bat species with ACE2 supporting the entry of indicated sarbecoviruses are analyzed (Extended Data Fig. 7a). Although PRD-0038 has been proposed as a sarbecovirus with broad ACE2 usage^42^, this virus and three other clade 3 sarbecoviruses display a moderate breadth in our study (Extended Data Fig. 7b). The RBD binding of the five sarbecoviruses with their optimal ACE2 orthologues, most are from their hosts, was further demonstrated through flow cytometry and Bio-layer interferometry (BLI) (Fig. 2c-d).

Notably, two close-related RBM type-I sarbecoviruses, GX-P2V and RaTG13, displayed contrasting ranges of ACE2 tropism (Fig.2a, b). Pseudovirus entry and RBD binding data based on swap mutants between the four residues on positions 493, 498, 501, and 505 (SARS-CoV-2 numbering) highlighted the critical role of position 501 residues in determining the breadth of ACE2 tropism (Fig. 2e-h, Extended Data Fig. 8a)^1,33,46,47^. Residue usage analysis of the six ACE2 positions (38, 41, 42, 353, 354, and 355) that are spatially close to position 501 (SARS-CoV-2 numbering) underscore an overall negatively charged surface among the 56 orthologues, thereby disfavoring D501 due to electrostatic repulsion (Fig. 2i). This hypothesis is further confirmed by similar phenotype of SARS-CoV-1, SARS-CoV-2, and RshSTT200 carrying D/T mutations at the same position (Extended Data Fig. 8b-c and Extended Data Fig. 9a-d). Since N501Y became dominant during SARS-CoV-2 spreading in humans, we also compared the multi-species ACE2 usage spectra of SARS-CoV-1 and SARS-CoV-2 carrying N or Y at position 501_SARS-CoV-248_. The result showed the Y mutation in this position resulted in reduced ACE2 tropism of SARS-CoV-1 but an expanded tropism in SARS-CoV-2 (Extended Data Fig. 9a-d). Structural analysis shows Y487_SARS-CoV-1_ may result in steric hindrance with local Y41_hACE2_ and K353_hACE2_, whereas the Y501_SARS-CoV-2_ instead forms a π-π stacking interaction with Y41_hACE2_, highlighting a virus-specific influence (Extended Data Fig. 9e-f)^49^.

The different PSV entry efficiencies of SARS-CoV-2, SARS-CoV-2-N501Y, and SARS-CoV-2- Omicron BA.1 in using different ACE2 orthologues indicated the presence of other residues affecting the ACE2 tropism of Omicron BA.1 other than the 501 residues, which is further confirmed by the authentic SARS-CoV-2 infection assays (Fig. 2j and Extended Data Fig. 10a). Fine mapping of the mutations in BA.1 underscores the critical contribution of residues 493 in impacting multi-species ACE2 tropism (Fig. 2j-k, Extended Data Fig. 10b-f).

Collectively, these data indicated that the overall multi-species ACE2 adaptiveness and species-specific ACE2 usage are affected by the RBM indel types and critical RBM residue usage, particularly at positions 501_SARS-CoV-2_ and 493_SARS-CoV-2_.

### The impact of RBM indels on ACE2 recognition

To investigate the impact of loop deletions on multi-species ACE2 adaptiveness, we generated chimeras with specific loop substitutions in Region 1 and 2. These comprise SARS-CoV-2 with single deletions in either Region 1 or 2 and other sarbecoviruses carrying partial or entire loop substitutions with SARS-CoV-2 equivalent sequences (Fig. 3a). The cellular expression and VSV package efficiency of all spike chimeras were validated by Western blot (Fig. 3b).

**Fig. 3.**
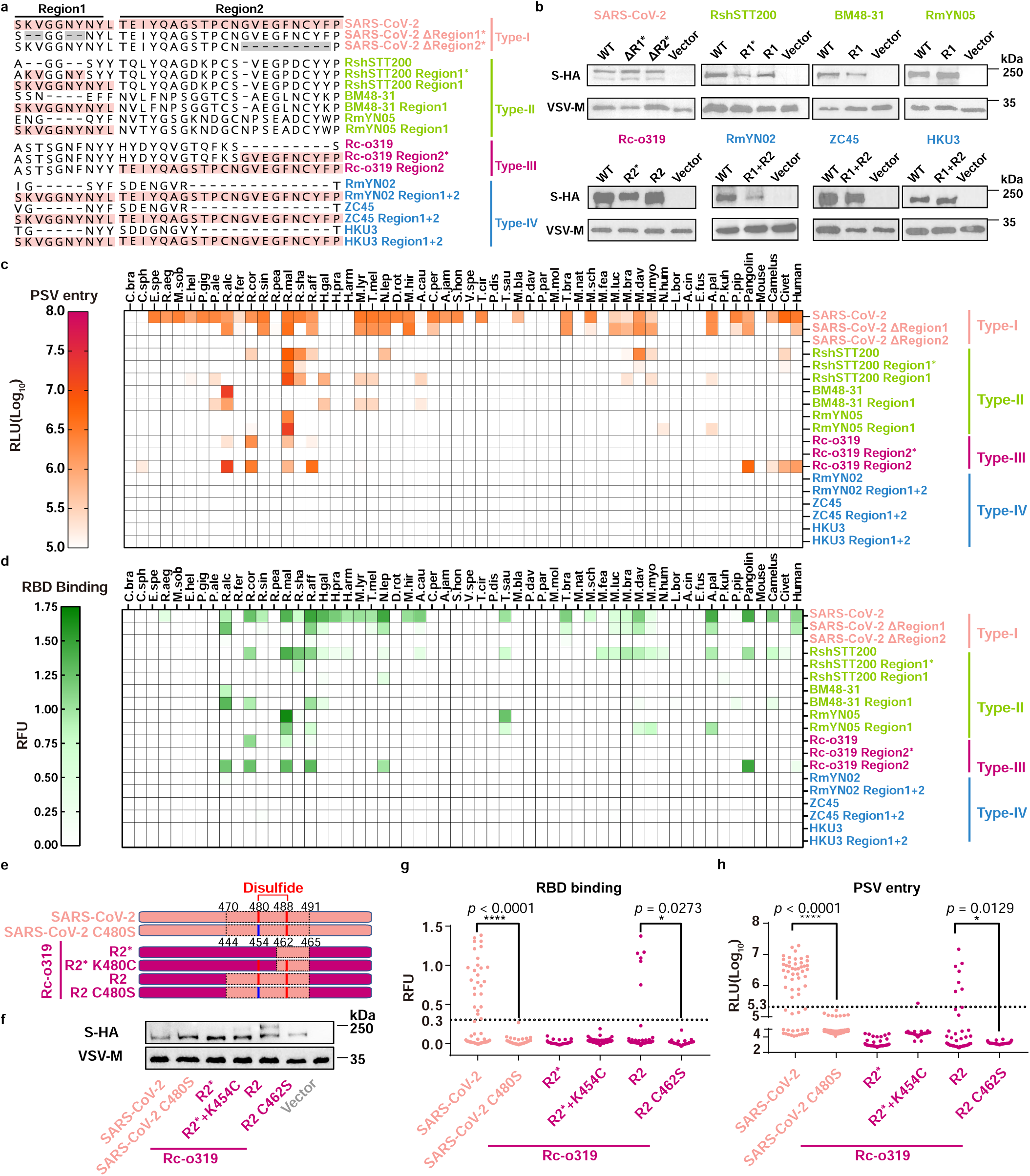
The impact of Region1 and Regon2 deletions on multi-species ACE2 tropism of different Sarbecoviruses. **a,** Illustration displaying the Region1 and Region 2 substitutions in sarbecoviruses of different indel types. Light pink: insertion corresponding to SARS-CoV-2 counterparts; gray: deletions corresponding to type II and type III sarbecoviruses. **b,** Western blot detecting the spike protein packaging efficiency in PSV particles. **c, d,** PSV entry **(c)** and RBD binding**(d)** efficiency of sarbecoviruses and their region substitution mutants in HEK293T stably expressing the 56 ACE2 orthologues. **e-h,** Disulfide bonds in Region2 is critical for multi-species ACE2 usage for sarbecoviruses. Cartoon illustration displaying the Region2 disulfide-related mutants based on SARS-CoV-2, Rc-o319 and their region 2 substitution mutants (**e**). Spike protein package efficiency (**f**), RBD binding efficiency (**g**), and PSV entry efficiency (**h**) of SARS-CoV-2 and Rc-o319 mutants in HEK293T stably expressing the 57 ACE2 orthologues. Dots represent different ACE2 orthologues. Dashed lines: background cut-off of RBD binding and PSV entry assays. Data are presented in c and d for n=2-3 biologically independent cells. Chi-squared test was used for statistical analysis of significance for g and h. *:P<0.05, ****:P<0.0001. RLU: Relative Luminescence units; RFU: Relative Fluorescence Units.

SARS-CoV-2 with a Region 1 4aa KVNY deletion (ΔRegion1*, relative to RshSTT200) displayed reduced multi-species ACE2 adaptiveness but still retained the capacity to use ACE2 from many species, including humans (Fig. 3c, d). However, the 9aa partial deletion in Region 2 (ΔRegion2*, relative to Rc-o319) completely abolished its ability to use all tested ACE2 orthologues, including R.cor ACE2. For BM48-31 and Rc-o19, region substitution by SARS-CoV-2 counterparts slightly increases the number of supportive ACE2 orthologues, yet still unable to achieve a broad tropism as RBM type-I sarbecoviruses. Indeed, multi-species ACE2 tropism can be reduced if unfavorable residues are presented in the loops with complemented length, as observed in RshSTT200 and Rc-o319. Thus, entire region 1 substitution rather than just filling-up the gaps (mutants marked with *) is sometimes necessary for maintaining or expanding multi-species ACE2 tropism, indicating the side chain of the loop is crucial in addition to the loop length (Fig. 3c, d). Notably, the highly conserved disulfide bridge is present in Rc-o319-R2 but not in Rc-o319-R2*, and its importance was further verified by the loss of ability of SARS-CoV-2 and Rc-o319-R2 with a C480S mutation^45,50^. However, introducing a disulfide to Rc-o319-R2* via K480C mutation remains insufficient to restore its ability to use ACE2, suggesting the presence of incompatible residues for Region 2 ACE2 interaction (Fig3. e-h).

Unexpectedly, substituting both regions (R1+R2) in the three RBM type-IV (clade 2) sarbecoviruses (ZC45, RmYN02, HKU3) failed to recover any detectable ACE2 usage in both binding and entry assays (Fig3. c, d). These data indicate the presence of determinants other than loop deletions that restrict ACE2 usage in RBM type-IV sarbecoviruses^3^.

### Clade-specific residues restricting ACE2 usage

We unexpectedly found that HKU3 and ZC45 remained unable to bind any ACE2 even with the entire RBM (aa439-508) replaced, indicating the presence of determinants restricting ACE2 recognition outside the RBM region (Extended Data Fig.11a). When comparing RBD sequences from 172 RBM type-IV (clade 2) sarbecoviruses with the other 96 ACE2-using sarbecoviruses, approximately twenty of clade 2 specific residues situated within or outside the RBMs were identified (Fig. 4a). It has been proposed that two residues (D496 and P502) within the Region 3 of RBM type-IV sarbecoviruses may restrict potential ACE2 interaction based on structural modeling, while the impact of this two residues, as well as other RBD residues, to ACE2 recognition remains to be investigated by cell-based functional assays^51^.

**Fig. 4.**
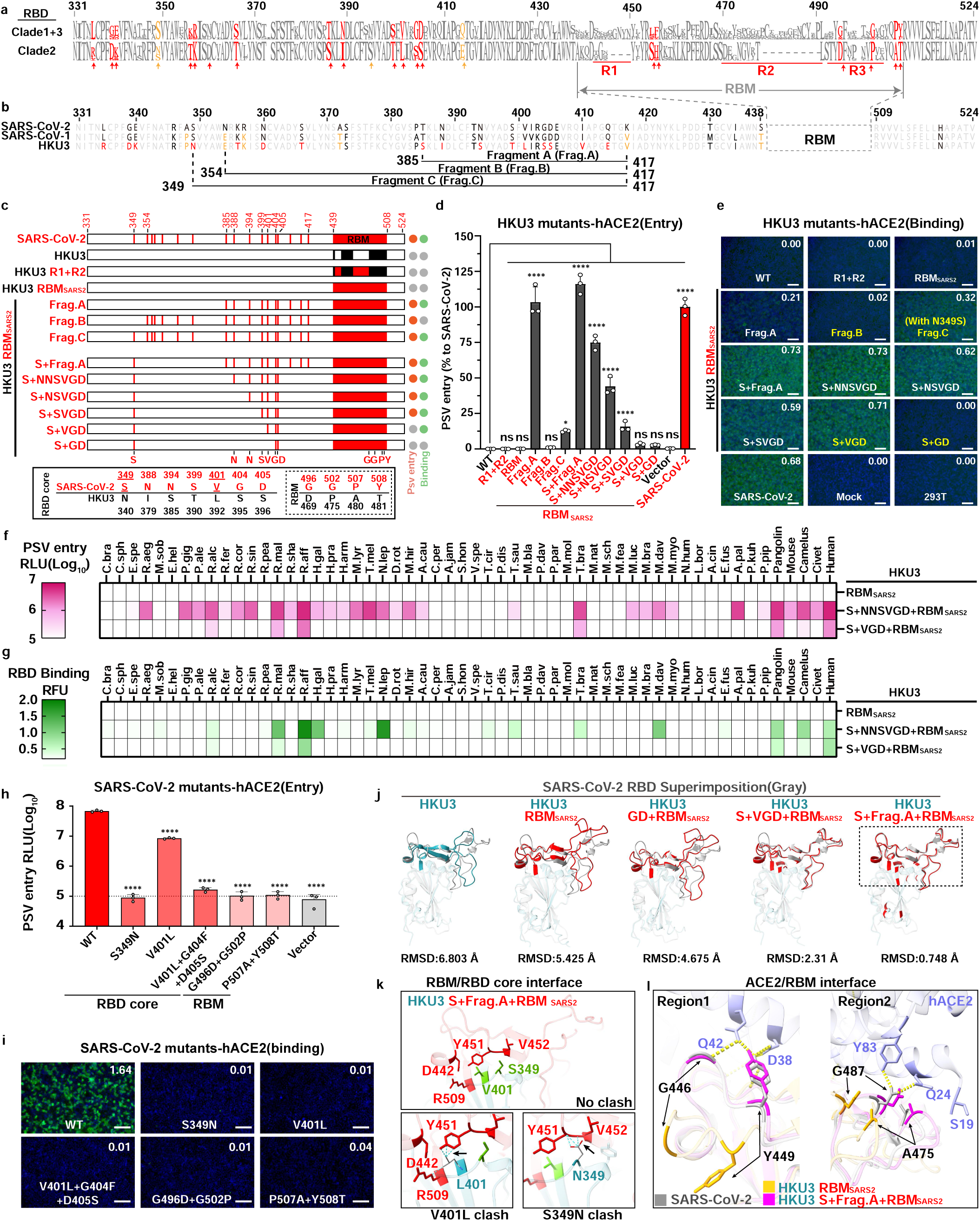
Fine mapping of Clade-2 specific residues outside the RBM restricting ACE2 recognition. **a,** RBD residue usage (SARS-CoV-2 numbering) of sarbecoviruses grouped by ACE2 usage. Red: Clade 2-specific residues. Orange: limited Clade 2 specificity. **b,** RBD sequences alignment of SARS-CoV-2, SARS-CoV-1 and HKU3. Red: HKU3-specific; Orange: shared by HKU3 and SARS-CoV-1 only. The three fragments (Frag.) for subsequent mapping are indicated. **c-g,** Fine mapping of residues restricting ACE2 usage outside the RBM. Mapping strategy for narrowing down the determinants critical for ACE2 recognition **(c)**. Orange and green circles: capability of using hACE2 for entry (>1% of SARS-CoV-2 entry) and binding (RFU>0.2), respectively. Gray: unable to use hACE2. Underlines: two critical residues dictating RBD binding. PSV entry **(d)** and RBD binding (**e**) of the HKU3 mutants carrying SARS-CoV-2 corresponding sequences in HEK293T-hACE2. Yellow highlighted the mutants critical for analyzing ACE2-restricting determinants. PSV entry (**f**) and RBD binding (**g**) of HKU3 mutants with restored ACE2 binding affinity. **h-i,** PSV entry **(h)** and RBD binding **(i)** of SARS-CoV-2 mutants carrying Clade 2-specific restricting residues in HEK293T-hACE2. **j-l**: Mechanisms of ACE2 restriction by clade-specific residues outside the RBM. RBD superimposition (**j**) of SARS-CoV-2 (PDB:6M0J) with HKU3 and HKU3-derived mutants predicted by Alphafold2. Red: SARS-CoV-2 equivalent sequences. RMSD: root mean square deviation based on RBM (69 Cα atoms). Close-up view (**k**) of V401L and S349N on HKU3 S+Frag.A+RBM mutations that form steric hindrance with RBM that potentially inducing the RBM conformational shift. The loss of ACE2 interactions in HKU3 carrying SARS-CoV-2 RBM compared with HKU3 S+Frag.A+RBM mutant (**l**). Blue dashed line: Clashes; Yellow dashed line: H-Bond. One-way ANOVA analysis, followed by Dunnett’s test for **d** and **h,** mean ± SD. Scale bar, 200 μm. Data representative of 2 or 3 independent experiments for d, e, h and i (n=3 biological replicates).

To identify the determinants restricting ACE2 recognition, we conducted RBM sequence swap analyses based on SARS-CoV-2 and HKU3, the sarbecoviruses showing the highest RBD protein identity (63.24%) in our study (Extended Data Fig. 2b). Sequence alignment of SARS-CoV-1, SARS-CoV-2 and HKU3 RBD displayed 16 HKU3-specific residues upstream of the RBM region (Fig. 4b). The dissection started from large fragment swaps and then proceeded to fine mapping of single residues (Fig. 4b, c). In addition to SARS-CoV-2 RBM (HKU3-RBM_SARS2_) replacement, further substituting fragment A (aa385-417) enabled HKU3 to use hACE2 for efficient entry but remained deficient in binding hACE2. Further extension by fragment B (aa354-417) and fragment C (aa349-417) underscore the critical contribution of S349_SARS-CoV-2_ for efficient binding (Fig. 4d, e). Fine mapping of fragment A highlighted the crucial role of six residues in position 388, 394, 399, 401, 404, 405 (SARS-CoV-2 numbering) that restricting hACE2 usage, all of which are clade-2 specific residues (Fig. 4c-e and Extended Data Fig.11b). The multi-species ACE2 usage spectra of HKU3-RBM_SARS2_ carrying S+NNSVGD, S+VGD mutations were demonstrated with improved ACE2 adaptiveness (Fig.4f, g). Similar results were obtained when testing another RBM type-IV sarbecovirus, ZC45 (Extended Data Fig. 12).

The restrictive effect of these clade 2-specific residues was further demonstrated by the loss of ACE2 usage of SARS-CoV-2 mutants. SARS-CoV-2 carrying the corresponding mutants within or outside the RBM region (S349N, V401L, V401L+G404S+D405S, G496D+G502P, and P507A+Y508T) all displayed a significantly reduced efficiency in use hACE2 (Fig. 4h, i). Structural modeling by AlphaFold2 suggests that HKU3 RBD carrying increasing substitutions of SARS-CoV-2 equivalent sequences resulted in a gradually decreased root mean square deviation (RMSD) when superimposing with SARS-CoV-2 RBD. The RMSD reduction is apparently due to the RBM conformational shift, indicating an RBM remodeling toward a structure more compatible with ACE2-binding (Fig. 4j).

Interestingly, the two clade-2 specific residues crucial for ACE2 binding, S349_SARS-CoV-2_ and V401_SARS-CoV-2_, are situated underneath the canonical RBM region. V401L and S349N (SARS-CoV-2 numbering) in HKU3 may interfere with the RBM conformation due to their relatively larger side chains (Fig. 4k). The resulting conformational shift may lead to the mismatch of critical residues for ACE2 interaction, thereby restricting ACE2 usage even with SARS-CoV-2 RBM sequences (Fig. 4l). Thus, it is very unlikely for so far identified clade 2 sarbecoviruses to gain ACE2 usage simply through RBM indels unless the entire RBD was substituted.

### Coevolution of RBM indels and ACE2 adaptiveness

To trace the coevolution of sarbecoviruses RBM indels and ACE2 adaptation, the functional data based on multi-species ACE2 usage were integrated with analyses based on RBD clades, RBM types, and residue usages in the two featured regions (Fig. 5a-c and Extended Data Fig.13). The multi-species ACE2 adaptiveness and hACE2 usage reveal the overall spillover risks of sarbecoviruses from different clades and RBM indel types (Fig. 5a-b). The close scrutiny of the sequence features unveils an intriguing indel pattern in Region 1, characterized by one or two centrally located glycine (G), while the sequence features in Region 2 underscore the presence of a complex indel in this region rather than a straightforward 9aa or 13/14 deletion in RBM type III and IV sarbecoviruses, respectively (Fig. 5c and Extended Data Fig.13). Notably, a potential evolutionary trace of Region 1 insertion was identified by the likely duplication of NY/NF sequences on the right side of the G. These analyses provide valuable insights for deciphering the evolutionary trajectories of various sarbecoviruses.

**Fig. 5.**
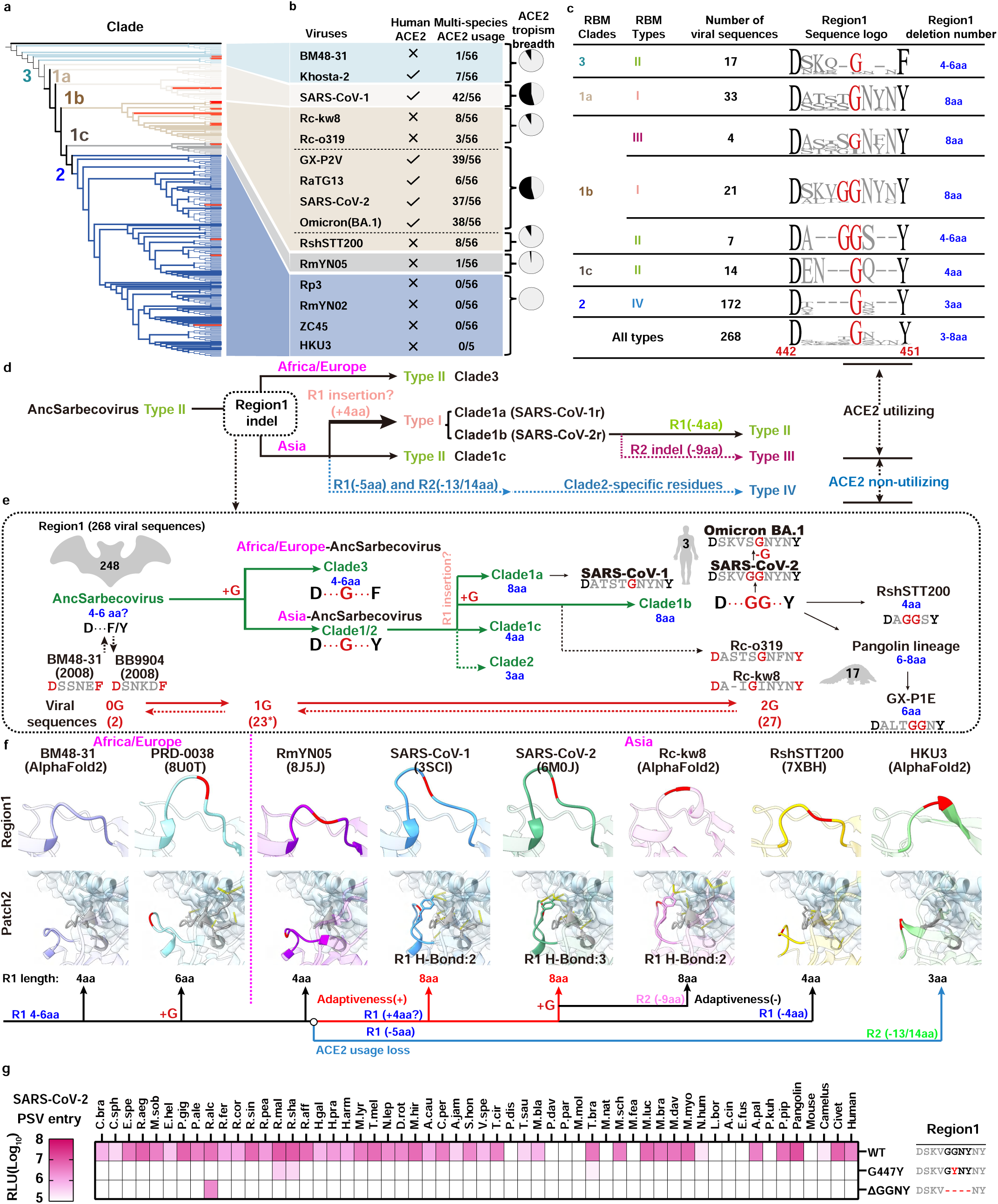
The proposed coevolution of sarbecovirus RBM indels and their impact on multi-species ACE2 adaptiveness. **a,** The phylogenetic tree based on RBD protein sequences using maximum likelihood analysis (details in Extended Data Fig. 1). The red lines mark the sarbecoviruses tested in this study. **b,** Summary of the number of supportive ACE2 orthologues (data based on Fig.2a and j with RLU>2×10^5^) and hACE2 compatibility of the indicated sarbecoviruses. Coloring is based on RBD clades. **c,** Region1 sequence logo (SARS-CoV-2 numbering) of sarbecoviruses grouped by different indel types in each clade. The highly conserved D442, F/Y451 for defining the boundary of Region 1 and the featured glycine (G) are highlighted in red. **d,** The proposed evolutionary of sarbecoviruses RBD clades and RBM indel types. **e,** Details of Region 1 sequence changes along with the formation of different clades during the evolution of sarbecoviruses in bats, pangolins, and humans. The Region 1 numbers in each group are indicated in blue. The emergence of the highly conserved glycine (1G) and the double G (2G) in most clade1b sarbecoviruses are highlighted in red. **f,** The RBD-ACE2 complex structures or models of sarbecoviruses with distinct Region 1 sequences. Region 1 is highlighted with distinct colors without transparency, the featured G is marked in red (upper). ACE2 and Region 3 is displayed in light blue and gray with transparency, respectively. Yellow dotted lines: H-bond or salt bridge. Dotted lines indicate the events with low confidence for d and **e**. **g**, PSV entry of SARS-CoV-2 R1 mutants in HEK293T expressing the indicated ACE2 orthologues.

Combining this information, we proposed an evolutionary pathway delineating the emergence of various RBM types, highlighted by critical events driving the evolution (Figure 5d). While the origin of the common ancestor of sarbecoviruses remains elusive, Africa/Europe sarbecoviruses (Clade 3) maintained a relatively ancient state of RBM indel type-II. The Asia sarbecoviruses underwent extensive evolution and developed into clade 1 and 2 sarbecoviruses with great genetic diversity. These viruses evolved in three different directions, each exhibiting distinct ACE2 adaptiveness. Clade 1c sarbecoviruses maintained RBM type-II with limited genetic diversities based on currently known sequences. Clade 2 sarbecoviruses underwent R1 (-5aa) and R2 (-13/14aa) indels and lost ACE2 usage, which is coupled with the emergence of clade 2-specific residues that further restricted ACE2 usage. On the other hand, clade 1a and 1b viruses underwent Region 1 insertion (or indels with increased residue numbers), generating the longest (8aa) Region 1 and superior multi-species ACE2 adaptiveness. Some Clade 1b viruses subsequently underwent further indels in Region 1 (-2-4aa) and Region 2 (- 9aa), which turned into RBM type II (e.g. RshTT200 and GX-P1E) and type III (e.g. Rc-o319 and Rc-kw8), respectively (Figure 5d).

While Region 1 is shorter than Region 2, it displayed more dynamic sequence changes in ACE2 using sarbecoviruses, fine-tuning the species-specific ACE2 adaptiveness. By contrast, no further RBM indel was observed in the 172 RBM type-IV (clade 2) sarbecoviruses, indicating that Region 1 may no longer be a crucial determinant for the adaptation of their receptor that other than ACE2. Intriguingly, despite the high variability in Region 1 sequences, only two out of the 268 sequences, BM48-31 and BB9904, lack a G in this region. In RBM type I sarbecoviruses with 8aa length, most clade 1b viruses maintained a double G (2G), whereas most 1a viruses kept an SG or TG in the middle (Figure 5c, e).

From a structural aspect, indels in Region 1 resulted in different loop lengths, with a G close to the turn of the loop. Interestingly, the longer Region 1 loop allows a closer distance and potential H-bond formation with ACE2, reinforcing ACE2 interactions along with the Region 3 loop, highlighting the importance of Region 1 length and its residue usage in multi-species ACE2 adaptiveness (Fig.5f). Our hypothesis is supported by the data showing a dramatic decrease of ACE2 fitness in SARS-CoV-2 carrying a G447Y mutation (Fig.5g and Extended Data Fig.14). Additionally, we observed a reduced multi-species ACE2 adaptiveness of the “G-free” 4aa deletion mutant, SARS-CoV-2-ΔGGNY (Region 1: DSKVNY), compared to SARS-CoV-2-ΔKVNY (Region 1: DSGGNY) (as shown in Figure 3), the former only recognize R.alc ACE2, similar to the phenotype of BM48-31 which also lacks G in region 1 (Fig.5g and Extended Data Fig.14). SARS-CoV-2-ΔGGNY may employ a similar ACE2 recognition mode as BM48-31, considering the importance of position 31_R.alc ACE2_ for both viruses in a swap mutagenesis experiment based on R.alc ACE2 and its closest orthologue, R.fer ACE2 (Extended Data Fig. 15).

## Discussion

The long-term and constant evolution of sarbecoviruses in rhinolophus bats drives the emergence of sarbecovirus clades with diversified RBM sequences. The frequent sequence changes, particularly indels, within the RBM pose challenges in predicting the potential of sarbecoviruses to cross species barriers and spill over to humans. To more precisely investigate the influence of indels and other critical residues on multi-species ACE2 usage, we propose a novel RBM indel-based classification, categorizing all currently identified sarbecoviruses into four distinct RBM indel types.

Our functional data, combined with extensive sequence analyses, led to a detailed summary of the ACE2 usage adaptiveness of sarbecoviruses within specific clades and RBM indel types (Extended Data Fig.16, Graphic Abstract). Despite with narrower ACE2 tropism, all tested sarbecoviruses carrying single deletions exhibited confirmed ACE2 usage, typically adapting well to ACE2 from their hosts. Furthermore, we proposed a hypothesis outlining the evolutionary history of sarbecoviruses exhibiting distinct RBM indel types. Since the number of sarbecovirus sequences from different clades is constantly increasing, the more intricate evolutionary history of sarbecoviruses remains to be updated or amended.

The driving force for the emergence of different sarbecovirus Region 1 and Region 2 indels remains elusive. Virus recombination may play a crucial role, considering that the RBM or even Region 1 has been predicted as a breaking point for combinations between sarbecoviruses^35^. Additionally, although various NTD-indels emerged in SARS-CoV-2 during the pandemic, so far, no indels have been detected in RBM Region 1 or Region 2, suggesting a different evolution mechanism of RBM indels formation of various sarbecoviruses in bats compared with that of SARS-CoV-2 in humans^52^.

Our results reveal a coevolution between sarbecovirus indels and multi-species ACE2 adaptiveness. Remarkably, the fine-tuning of RBM Region 1 through various indels and specific side chains promotes the emergence of various sarbecoviruses with distinct multi-species ACE2 usage spectra. This could be attributed to the dispensable nature of Region 1 for interaction with a specific ACE2 orthologue due to the compensation of the Region 3 loop without indels, while additional interactions mediated by the extended Region 1 loop in RBM type-I sarbecoviruses might be crucial for achieving better multi-species ACE2 adaptiveness to facilitate host jumping. Interestingly, a conserved G within Region 1 suggests that better flexibility or the absence of a large side-chain in this region may confer some evolutionary advantage. Comparatively, indels in Region 2 are less diversified than in Region 1 and generally have a more dramatic impact on ACE2 recognition than deletion in Region 1, or even result in the switch of ACE2 usage to another yet-unknown receptor. Notably, the deficiency in multi-species ACE2 adaptiveness does not mean a lack of ability to use hACE2, as is exemplified in Khosta-2 or other Clade 3 sarbecoviruses mutants ^1,42–44^.

Filling RBM deletions with SARS-CoV-2 counterparts does not guarantee a broader ACE2 usage spectrum, sometimes instead resulting in reduced or lost ACE2 usage. This underscores the enhanced ACE2 adaptiveness achieved during adaptive evolution, with both length and residues being optimized in specific indels. Consequently, substituting the entire loop sometimes is necessary for achieving higher ACE2 compatibility. However, RBM type-IV sarbecoviruses, even after gap-filling or entire RBM substitution, remained unable to use any ACE2 orthologues, which led to the identification of clade-specific determinants outside the RBM that restrict ACE2 usage, probably a consequence of adaptation to another receptor usage. Interestingly, some critical RBD core residues are underneath the RBM, indirectly restricting ACE2 binding by affecting the RBM conformation. Future structural analysis could shed light on the molecular details of how these determinants affect receptor recognition.

It should be noted that although ACE2 fitness serves as the primary barrier for sarbecoviruses to cross species, ACE2 compatibility alone does not guarantee susceptibility at the animal level. Other factors, such as host protease, immune response, and viral replication efficiency, also affect host tropism, which can be verified by authentic viruses and in vivo studies in the future^11,53,54^.

In conclusion, our RBM indel type classification offers a more precise way to describe sarbecoviruses when integrated with RBD phylogenetic information. Our functional ACE2 usage data elucidate the underlying mechanism governing multi-species ACE2 usage and adaptiveness, shaped by multiple factors such as the presence and features of RBM loop deletions, RBM disulfide bridges, critical RBM residues for direct interaction, and ACE2 usage restricting residues within and outside the RBM. These findings establish a solid scientific foundation for risk assessment and viral surveillance to mitigate potential future zoonoses caused by these viruses.

## Author contributions

Conceptualization, H.Y., and J.Y.S; methodology, J.Y.S, Y.M.C, M.X.G, Y.H.S, C.L.W, C.L, C.B.M, P. L, Q.X, L.L.S, F.T, M.L.H, X.Y, X.Y, Z.X.M, Y.C.S; data analysis, J.Y.S, Y.M.C, M.X.G, Y.H.S; writing—original draft, H.Y., J.Y.S, Y.M.C; writing—review & editing, H.Y., J.Y.S, Z.L.S.; supervision and funding acquisition, H.Y..

## Acknowledgments

We are grateful to the funding support from National Key R&D Program of China (2023YFC2605500 to H.Y.), National Natural Science Foundation of China (NSFC) projects (82322041, 32270164, 32070160 to H.Y.), Fundamental Research Funds for the Central Universities (2042023kf0191, 2042022kf1188 to H.Y.), and Natural Science Foundation of Hubei Province (2023AFA015 to H.Y.). We thank Huabin Zhao (Wuhan University) for his help in providing the coding sequences of many bat ACE2 orthologues. We thank Ming Guo (Wuhan University) for his help in conducting SARS-CoV-2 authentic viruses related experiments in ABSL-3. We thank Qiang Ding (Tsinghua University) for his kind offer of several plasmids expressing mammalian ACE2 orthologues. We also want to express our gratitude to the core facilities and ABSL-3 laboratory of the Key Laboratory of Virology, Wuhan University.

## Data and code availability

This study did not generate custom computer code.

Any additional information required to reanalyze the data reported in this paper is available from the lead contact upon request.

## RESOURCE AVAILABILITY

### Lead contact

Further information and requests for resources and reagents should be directed to and will be fulfilled by the lead contact, Huan Yan

### Materials availability

All reagents generated in this study are available from the lead contact with a completed Materials Transfer Agreement.

## Materials and Methods

### Cell culture

HEK293T cells (ATCC, CRL-1586) and their derivatives were maintained in Dulbecco’s modified eagle medium (DMEM; Gibco) supplemented with 10% fetal bovine serum (FBS), 2.0mM L-Glutamine, 110 mg/L sodium pyruvate, and 4.5 g/L D-glucose. I1-Hybridoma (CRL-2700), secreting a monoclonal antibody targeting VSV glycoprotein (VSV-G), was maintained in Minimum Essential Medium with Earlès salts and 2.0 mM L-Glutamine (MEM; Gibco). All cells were cultured at 37℃ in 5% CO_2_ with regular passage every 2-3 days.

### Gene sequences

Sarbecovirus spike sequences are retrieved from NCBI Virus and GISAID databases. The keywords used for search include "Betacoronavirus," "Sequence length between 1000-1400" "protein" and "NOT Homo sapiens" for NCBI and "bat," "pangolin," "civet," coronaviruses for GISAID. A comprehensive collection of 2318 Betacoronavirus spike protein sequences was obtained. After extracting 876 sarbecovirus sequences through phylogenetic analysis using Geneious, the dataset was refined to 268 unique sequences for further analysis by excluding redundant entries. The ACE2 orthologues sequences were summarized by previous reports^30^. Several additional ACE2 orthologues tested in this study include *Rhinolophine Malayanus* (Provided by Professor Huanbin Zhao, Wuhan University, China), *Rhinolophus shameli* (GenBank: MZ851782), *Rhinolophus cornutus* (GenBank: BCG67443.1), *Rhinolophus sinicus isolate Rs-3357*(GenBank: KC881004.1), *Rhinolophus affinis* (GenBank: QMQ39227.1), *Manis javanica (Pangolin)*(GenBank: XP_017505752.1), *Mouse* (GenBank: NP_001123985), *Camelus* (GenBank: XP_006194263), *Civet* (Protein: Q56NL1), *Rhinolophus alcyone* (Protein: ALJ94035.1). Human Aminopeptidase N precursor (APN) (Protein: NP_001141.2) was included as a negative control. The sources and accession numbers of the receptors and the 268 sarbecovirus were summarized in **Supplementary data S1**.

### Bioinformatic analysis

Amino acid or nucleotide sequences from viruses or ACE2 orthologues were aligned using Mafft v7.450^55^. Phylogenetic trees were generated with IQ-Tree (version 2.0.6)^56^ using a Maximum Likelihood model with 1000 bootstrap replicates. Tree annotations were performed using iTOL (https://itol.embl.de/). Sequence identities were analyzed by Geneious prism (https://www.geneious.com/) after aligned by Mafft. The residue usage frequency (Sequence Logo analysis) was generated by the Geneious Prime.

### Plasmids

The coding sequences of various coronavirus spikes and their derivatives were human codon optimized and cloned into the pCAGGS vector with a C-terminal 18-amino acids replaced with an HA tag (YPYDVPDYA) for improving VSV pseudotyping efficiency and enabling detection^57,58^. The plasmids for expressing ACE2 orthologues are constructed by inserting human codon-optimized ACE2 sequences into a lentiviral transfer vector (pLVX-IRES-puro) with a C-terminal 3×FLAG-tag (DYKDHD-G-DYKDHD-I-DYKDDDDK) for detection. The plasmids expressing the recombinant coronaviruses RBD human IgG Fc (RBD-hFc) fusion proteins were constructed by inserting RBD sequences corresponding to SARS-CoV-2 (aa331-524) containing an N-terminal CD5 secretion signal peptide (MPMGSLQPLATLYLLGMLVASVL) and a C-terminal hFc-twin-strep tandem tags for purification and detection.

### ACE2 stable expression cell lines

ACE2 stable expression cell lines were established as previously reported^30,59^. Briefly, lentivirus carrying the ACE2 genes was generated by co-transfecting pLVX-IRES-puro-ACE2 orthologues, pMD2G (plasmid no. 12259; Addgene), and psPAX2 (plasmid no. 12260; Addgene) into HEK293T cells. HEK293T cells were subsequent transduced with the lentiviruses, and the stable cells expressing ACE2 orthologues were selected in the presence of puromycin (1 μg/ml). The expression levels of ACE2 orthologues were assessed using an immunofluorescence assay as previously reported^30^. Briefly, HEK293T cells were fixed with 4% paraformaldehyde for 10 min at room temperature, permeabilized with 0.2% Triton X-100/PBS for 10 min, and blocked with 1% BSA for 30 min at 37 ℃. Subsequently, the cells were incubated with M2 antibody (anti-FLAG-tag, catalogue no. F1804A-5MG; Sigma) at 4 ℃ for 1 hour. After three washes with PBS, the cells were treated with 2 μg/ml Alexa Fluor 594- conjugated goat anti-rabbit IgG (catalogue no. A11032; Thermo Fisher Scientific). Nucleus were stained blue with Hoechst 33342 (1:5,000 dilution in PBS). Images were captured with a fluorescence microscope (MI52-N; Mshot). Relative fluorescence unit of Alexa Fluor 596 and Hoechst 33342 was quantified by Thermo Varioskan LUX. The expression of most ACE2 orthologues were also verified by Western Blot analysis in our previous reports^30^.

### Recombinant protein expression and purification

Recombinant RBD-hFc fusion proteins or ACE2 ectodomains (amino acid sequences 18-740 correspond to Human ACE2) fused with FLAG-strep-tag proteins were generated through transient transfection of HEK293T cells using Lipofectamine 2000. The transfected cells were cultured in SMM 293-TIS Expression Medium (Serum-free, without L-Glutamine) (Sino Biological). The supernatant, containing the recombinant proteins, was collected at 2, 4, and 6 days post-transfection, and the expression was confirmed through Western Blot analysis using the Goat Anti-Human IgG-Fc secondary Antibody (HRP) (SinoBiological Inc, SSA002) for RBD or the M2 antibody for ACE2. Protein purification was performed using Protein A/G Plus Agarose (Thermo Fisher Scientific) for RBD and Strep-Tactin®XT 4Flow® high capacity resin (IBA) for ACE2 ectodomains. The protein concentration was quantified using the BCA protein determination kit (EpiZyme) and SDS-PAGE with Coomassie blue staining was employed for analysis.

### Live cell RBD binding assay

HEK293T cells stably expressing ACE2 were seeded in poly-D-lysine-treated 96-well plates. After 12 hours, with cells were incubated with RBD-hFc protein (4 μg/ml) in growth medium for 0.5 hours at 4 ℃. Subsequently cells were washed with DMEM twice, and then treated with Alexa Fluor 488-conjugated goat anti-human IgG (catalogue no. A11013; Thermo Fisher Scientific) at a concentration of 2 μg/ml in DMEM with 2% FBS for 30 minutes (min) at 4 ℃. Hoechst 33342 (1:5,000 dilution in PBS) was utilized for nuclear staining. Following fixation with methanol, images were captured by fluorescence microscopy (MI52-N; Mshot), and the fluorescence intensity was analyzed using Thermo Varioskan LUX Alexa

### Flow cytometry

HEK293T cells stably expressing ACE2 orthologues (*R.aff, R.sha, R.alc, R.mal,* and *R.cor*) were cultured in 6-well plates for 12 hours. Cells were detached by 5mM EDTA and washed twice by PBS, and then incubated with indicated proteins (RaTG13 RBD, RshSTT200 RBD, BM48-31 WT RBD, BM48-31 A480Y RBD, RmYN05 RBD, Rc-o319 RBD with hFc tag) at a concentration of 20 μg/ml for 30 min at 4℃. Following three PBS washes, cells were stained with 488-conjugated goat anti-human IgG (1:1000, Alexa Fluor) for 30 min. Subsequently, flow cytometry analysis was performed using a CytoFLEX analyzer, collecting 10,000 events per sample. In a separate assay demonstrating the sensitivity of live cell binding assay, HEK293T cells expressing hACE2 were plated 12 hours before incubation with two-fold serial diluted SARS-CoV-2 RBD-hFc (from 20 μg/ml) for 30 min. After three PBS washes, cells were stained with 488-conjugated goat anti-human IgG (1:1000, Alexa Fluor) and subjected to Flow cytometry analysis. For the pseudoviruses entry assays, GFP expressing VSV pseudotypes was 10-fold serial diluted from 1×10^6^ TCID_50_/ml. After 12 hours post infection incubation, cells were washed with PBS and trypsinized for analysis. FlowJo V10 software was employed for data analysis.

### Biolayer interferometry

The Octet RED96 system (ForteBio, Menlo Park, CA) was employed to determine the apparent affinity (Kd, app, due to the potential dimerization or ACE2) between the RBD and ACE2. The buffer for analysis was phosphate buffer saline with 0.05% Tween20 (PBST). The RBD (10 μg/ml) was captured on ProA biosensors, followed by binding of ACE2 ectodomains at 2-fold serial dilutions ranging from 500 nM for 120s, followed by dissociated in the PBST for additional 300s. Kinetics was modeled in a 1:1 using ForteBio Octet analysis software v12.2.0.20 (ForteBio, Menlo Park, CA). Mean KD values were derived by averaging all binding curves that conformed to the theoretical fit with an R^2^ value≥ 0.95.

### Pseudovirus production and entry assays

Pseudovirus incorporating coronaviruses spike proteins were produced using a vesicular stomatitis virus (VSV)-based system with slight modifications to a well-established protocol^57,60,61^. In general, HEK293T cells were transfected with plasmids expressing S proteins through Lipofectamine 2000 (Biosharp, China). After 24 hours, the transfected cells were infected with VSV-dG-EGFP-FLuc (1×10^6^ TCID50/ml) diluted in DMEM followed a two-hour incubation on a shaker at 37 ℃, the cells were replenished with DMEM containing anti-VSV-G monoclonal antibody (I1, 1 μg/ml). After 24 hours, the pseudovirus-containing supernatant was harvested, clarified at 12,000 revolutions per minute (rpm) for 5 min at 4 ℃, and shored at -80 ℃. For the viral entry assay, the HEK293T cell lines expressing various ACE2 orthologues were inoculated with pseudotyped viruses in DMEM with 10% FBS. In general, 3×10^4^ trypsinized cells were incubated with pseudovirus (1.5×10^5^ TCID50/100 μL) in a 96-well plate to allow cell attachment and pseudovirus entry. At 16-20 hpi, images of the infected cells were captured by a fluorescence microscope (MI52-N; Mshot). Intracellular luciferase activity was determined using a Bright-Glo Luciferase Assay Kit (Promega Corporation, E2620) and measured with a Thermo Varioskan LUX, SpectraMax iD3 Multi-well Luminometer (Molecular Devices) or a GloMax 20/20 Luminometer (Promega Corporation).

### Authentic virus infection

The SARS-CoV-2 WT strain (IVCAS 6.7512) was provided by the National Virus Resource, Wuhan Institute of Virology, Chinese Academy of Sciences. The BA.1 strain (YJ20220223) was provided by Hubei Provincial Center for Disease Control and Prevention. SARS-CoV-2 authentic viruses related experiments were conducted in ABSL-3 facility at Wuhan University with the approval from the Biosafety Committee of ABSL-3 lab. HEK293T cells expressing ACE2 orthologues were seeded in poly-lysine-treated 96-well plates (1.25×10^5^ cells/well). After a 12 hours incubation period, SARS-CoV-2 strains (WT and Omicron BA.1) were introduced to different stable cells and incubated for 1-2 hours. Following a medium change to DMEM with 2% FBS, cells were cultured for 24 hours, fixed with methanol, and treated with anti-SARS-CoV-2 Nucleocapsid (N) antibody (catalogue no. 40143-MM05; Sino Biological) at 1:1000 for one hour at 37 ℃. After PBS wash, cells were treated with secondary antibody (Alexa Fluor 594) and Hoechst 33342 (1:10,000 dilution in PBS) for nuclei staining. Images were captured using a fluorescence microscope (MI52-N, Mshot, China).

### Structural analysis

Protein structures and complex were predicted by predicted by AlphaFold2 and HDOCK^62–64^. Briefly, AlphaFold2, implemented in ColabFold, was utilized with default settings for predicting the protein structures of various sarbecovirus RBDs and ACE2 orthologues. The top ranked model was used for all subsequent analyses. The docking of the ACE2 ectodomain in complex with RBD was accomplished using HDOCK (v.1.1). Structural representations and analyses were carried out within ChimeraX. The hydrogen bonds and clashes between the displayed amino acids were analyzed using the H-bonds and clashes command. RMSD values for structural superimpositions were calculated using the matchmaker command. The reported RMSD values specifically pertain to RBM Cα atoms. The following cryo-EM complex structures in the PDB database were also used for structural analysis in this study: human ACE2/SARS-CoV-2-RBD (Protein Data Bank 6M0J), human ACE2/SARS-CoV-1-RBD (3SCI), human ACE2/RaTG13-RBD (7DRV), human ACE2/GX-P2V-RBD (7DDP), human ACE2/SARS-CoV-2 alpha variant-RBD (7EKF), human ACE2/RshSTT200-RBD (7XBH), Rhinolophus alcyone ACE2/PRD-0038-RBD (8U0T), and RsYN04 RBD/antibody S43 (8J5J).

### Western Blot

To examine the intracellular sarbecoviruses spike protein expression levels, HEK293T cells were transfected with plasmids encoding the viral spike proteins fused with a C-terminal HA-tag. After 24 hours, cells were washed with PBS, lysed on ice for 10 min in 2% TritonX-100/PBS containing 1mM PMSF (Beyotime, ST506). The cell lysates were clarified by centrifugation at 12,000 rpm for 5 mins at 4 ℃. The supernatants were mixed with 1:5 (v/v) 5× SDS-loading buffer and incubated at 95 ℃ for 5 min. For evaluating the spike protein levels in pseudovirus (PSV) particles in the cultured medium, PSV was concentrated with a 30% sucrose cushion (30% sucrose, 15 mM Tris–HCl, 100 mM NaCl, 0.5 mM EDTA) at 20,000×g for 1.5 hours at 4 ℃. The concentrated PSV was then resuspended in 1×SDS loading buffer and incubated at 95 ℃ for 30 min. Following SDS-PAGE and PVDF membrane transfer, the blots were blocked with 10% milk in PBS containing 0.1% TBST (20 mM Tris-HCl pH 8.0, 150 mM NaCl) supplemented with 0.05% Tween-20 at room temperature for 1 hour. Primary antibodies targeting HA (MBL, MBL-M180-3), β-tubulin (Immunoway, YM3030), or VSV-M (Kerafast, EB0011) were applied at a 1:10,000 dilution in TBST with 1% milk. After three washes with TBST, blots were incubated with the secondary antibody Peroxidase AffiniPure Goat Anti-Mouse IgG (H+L) (Jackson Immuno Research, 115-035-003). Blots were further washed three times before chemiluminescence detection (SQ201, Yamei Biotech) using the ChemiDoc MP Imaging System (Bio-Rad).

### Geographical distribution of bat species

The global distribution data of bat species were obtained from the IUCN Red List of Threatened Species 2020, the base layer of the map (version 5.1.1) was sourced from Natural Earth, available at (https://www.naturalearthdata.com/downloads/110mcultural-vectors/). GeoScene Pro 21 was utilized to visualize and analyze the bat distribution data.

### Statistical analysis and data presentation

Most experiments were conducted 2-4 times with three biological repeats. Representative results were shown. Heat maps were generated based on RLU or RFU values, with background (control cells expressing APN) signals subtracted. Data are presented as means ± standard deviation(SD) as indicated in the figure legends. All statistical analyses were conducted using Prism 7 software (GraphPad). Two-tailed unpaired (Student’s) t-test was performed if only two conditions were compared. One-way ANOVA analysis, followed by Dunnett’s test, was employed for multiple comparisons. The association between the entry/binding efficiency and the presence of RBM disulfide was assessed using the chi-squared test. P < 0.05 was considered significant. *P < 0.05, **P < 0.01, ***P < 0.005, and ****P < 0.001.

---

**Extended Data Fig. 1.**
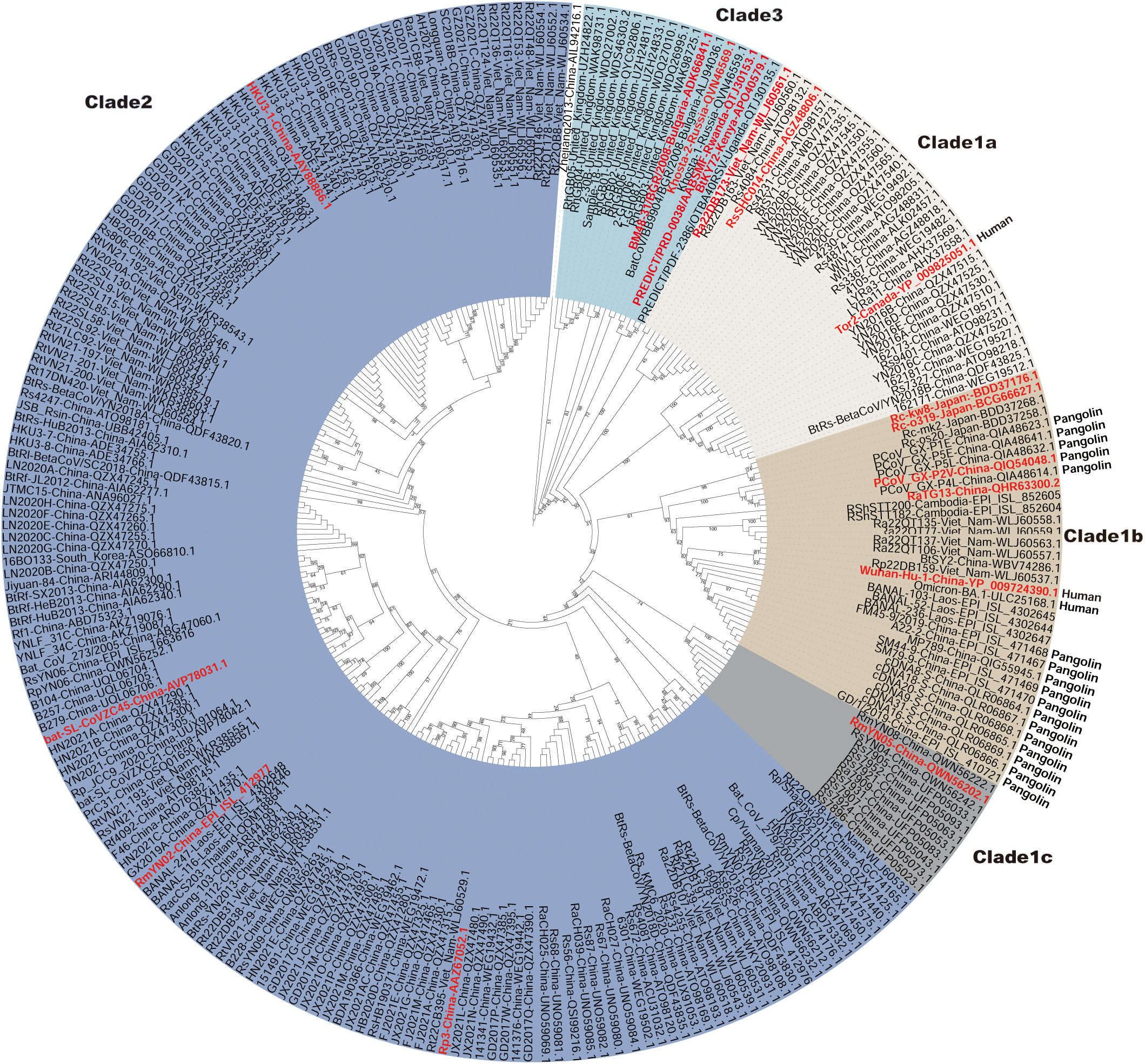
Phylogenetic tree based on 268 sarbecoviruses RBD amino acid sequences. The cladogram is generated by iq-tree through maximum likelihood analysis. Viruses selected in subsequent experiments are highlighted in red.

**Extended Data Fig. 2.**
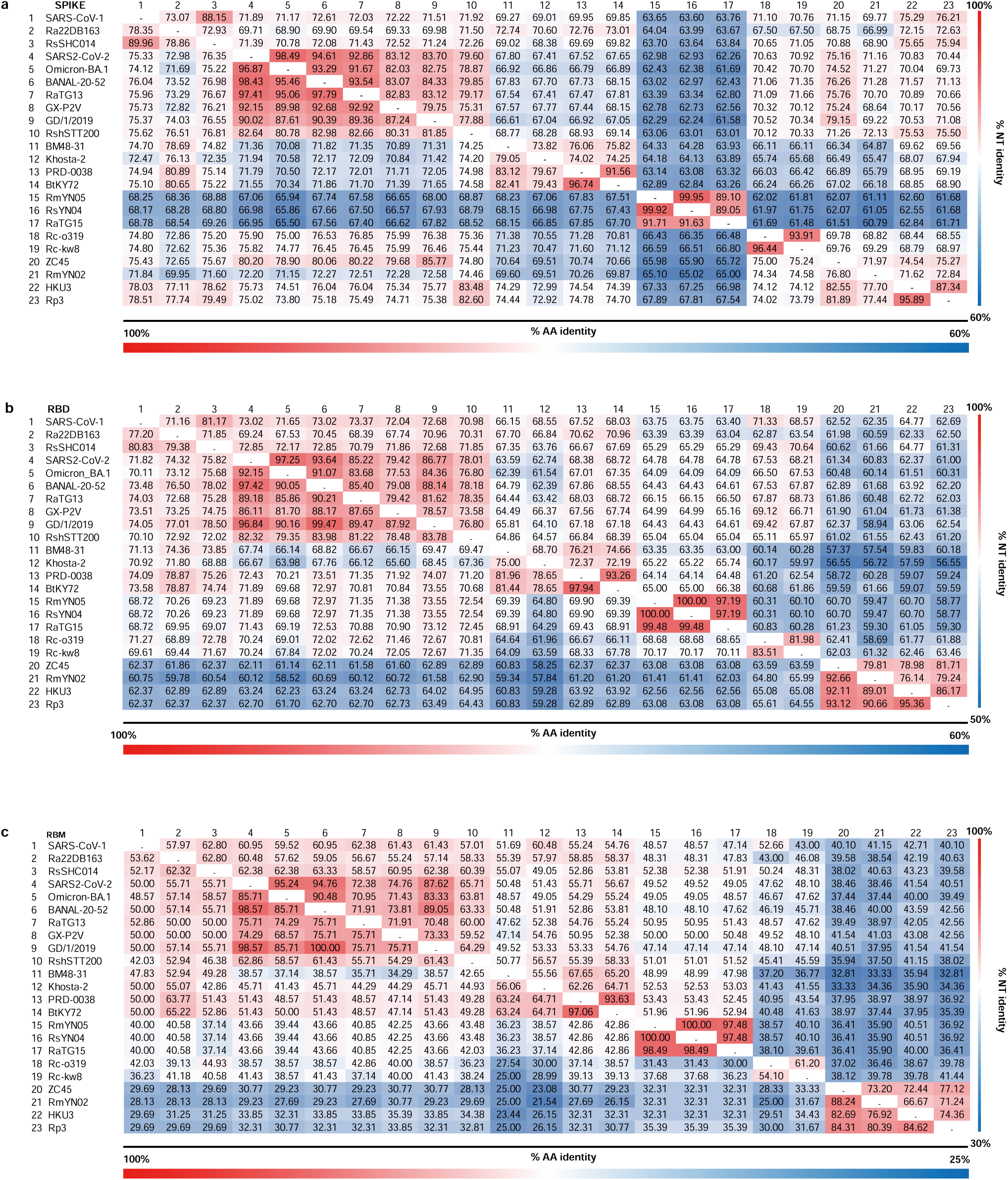
Protein and nucleotide identity matrices based on Spike (a), RBD (b) and RBM (c) sequences from 23 representative sarbecoviruses. The Nucleotide (NT) and amino-acid (AA) sequences are aligned by MAFFT, and the identities are analyzed by Geneious.

**Extended Data Fig. 3.**
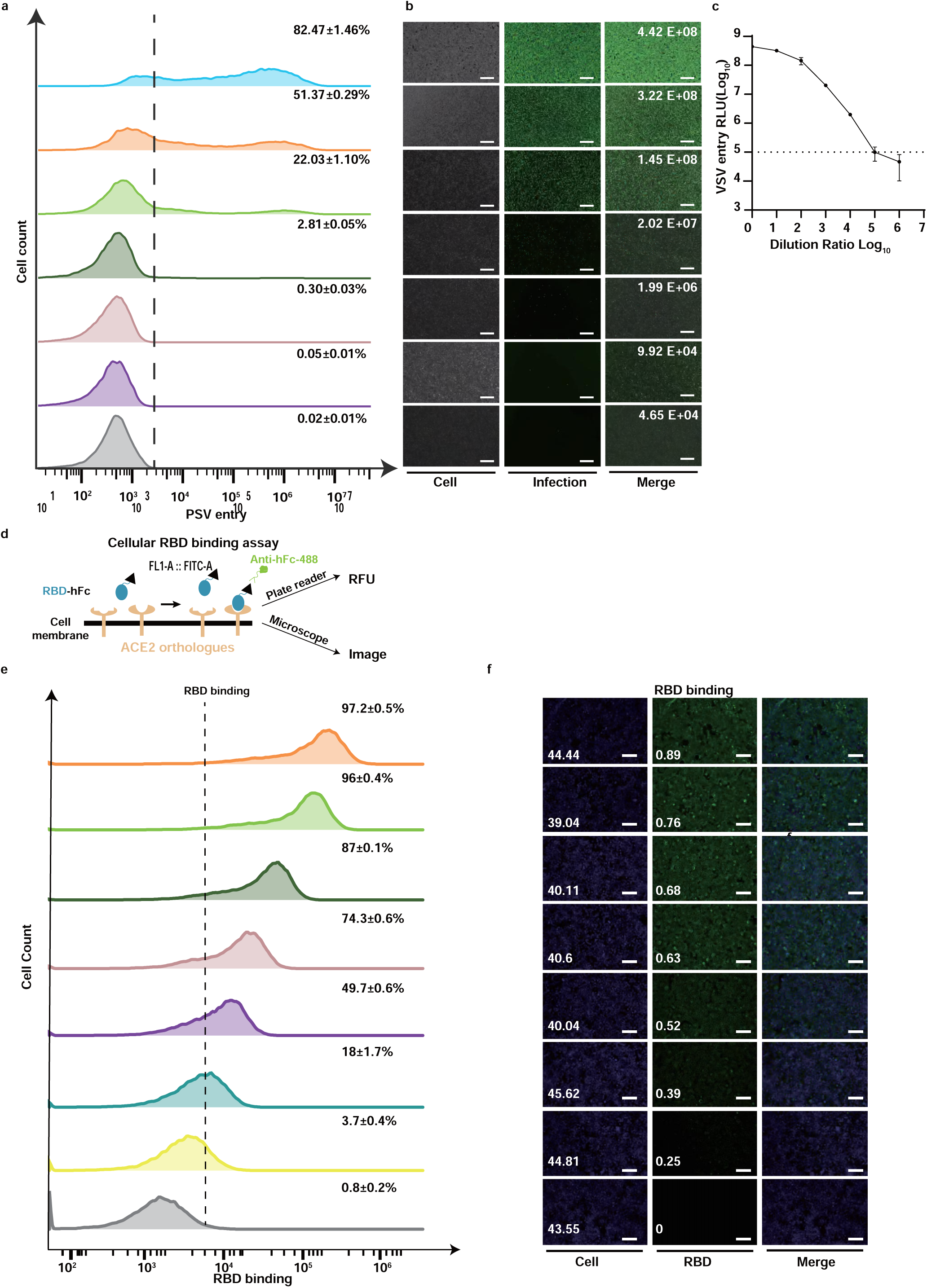
Characterization of quantitative RBD binding and PSV entry assays. **a-c,** Comparison of PSV entry efficiencies demonstrated by flow cytometry (**a**), GFP expression (**b**) and RLU (**c**). **d,** Schematic illustration of cellular RBD-hFc binding assay. **e, f,** Comparison of binding efficiencies based on flow cytometry (**e**) and quantitative immunofluorescence (**f**) assays. Scale bar, 200 μm.

**Extended Data Fig. 4.**
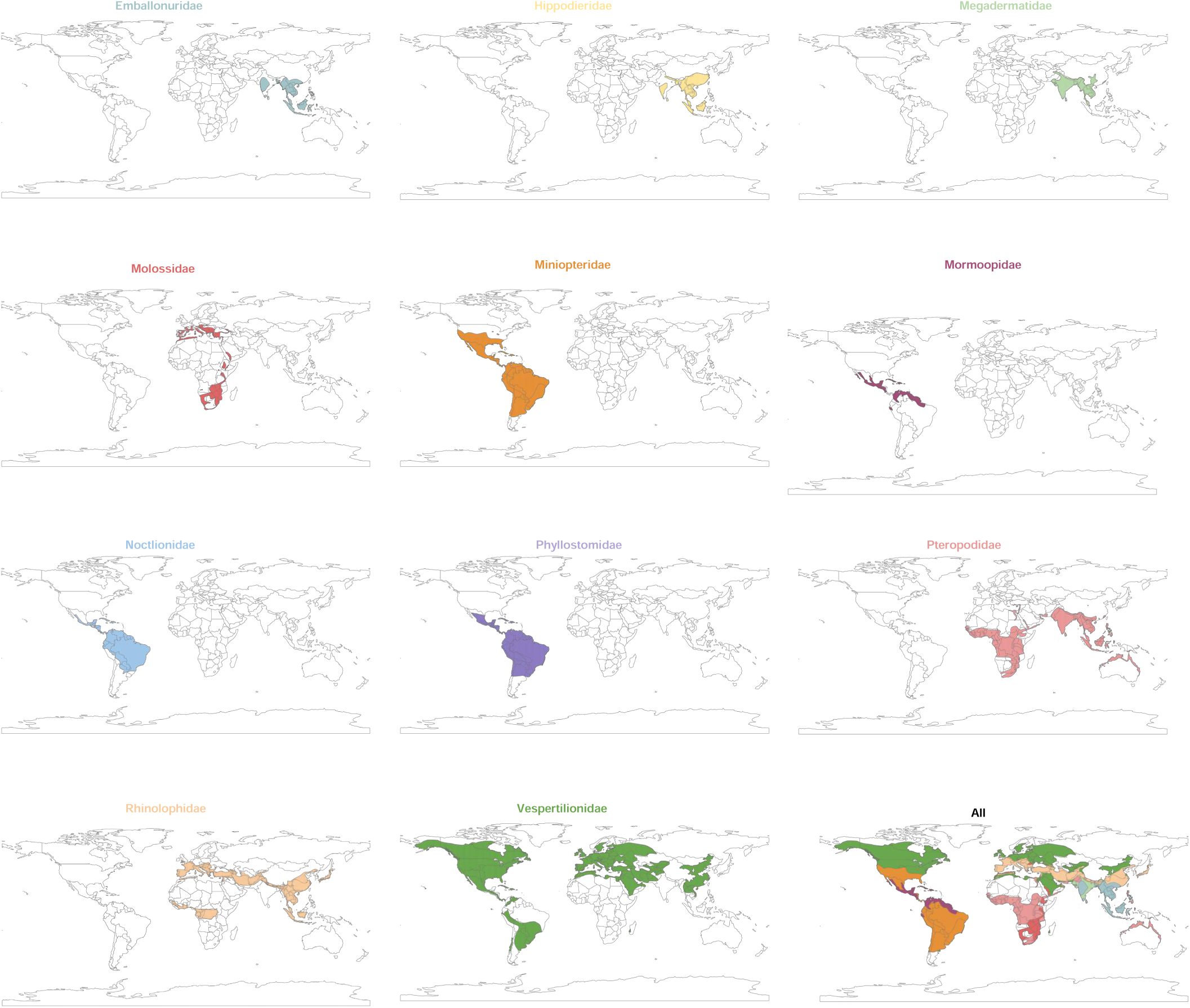
The geographical distribution of 11 bat families encompassing the 51 bat species in this study. Data are retrieved from The IUCN Red List website (https://www.iucnredlist.org/) and the distribution are generated by the GeoScene Pro software.

**Extended Data Fig. 5.**
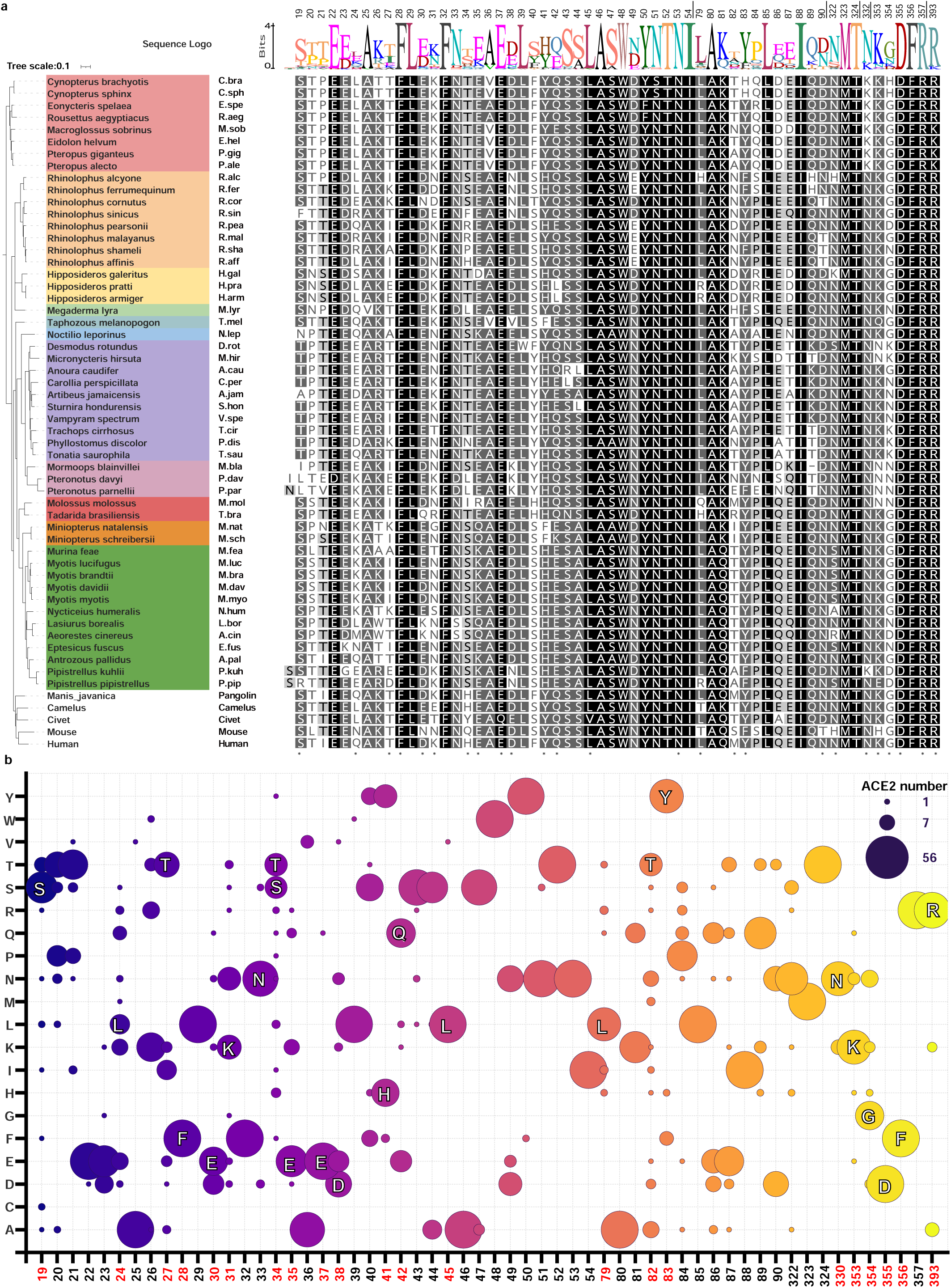
Phylogenetic and sequence analysis of the 56 ACE2 orthologues. **a,** The phylogenetic tree generated by iq-tree with maximum likelihood analysis. The multi-sequence alignment and sequence logo are analyzed by MAFFT and Geneious, respectively. Asterisks: critical residues for SARS-CoV-2 interaction (PDB: 6M0J). **b,** Bubble chart demonstrating the amino-acid usage of the viral binding sequences of 56 ACE2 orthologues. Residues critical for SARS-CoV-2 interaction are highlighted in red.

**Extended Data Fig. 6.**
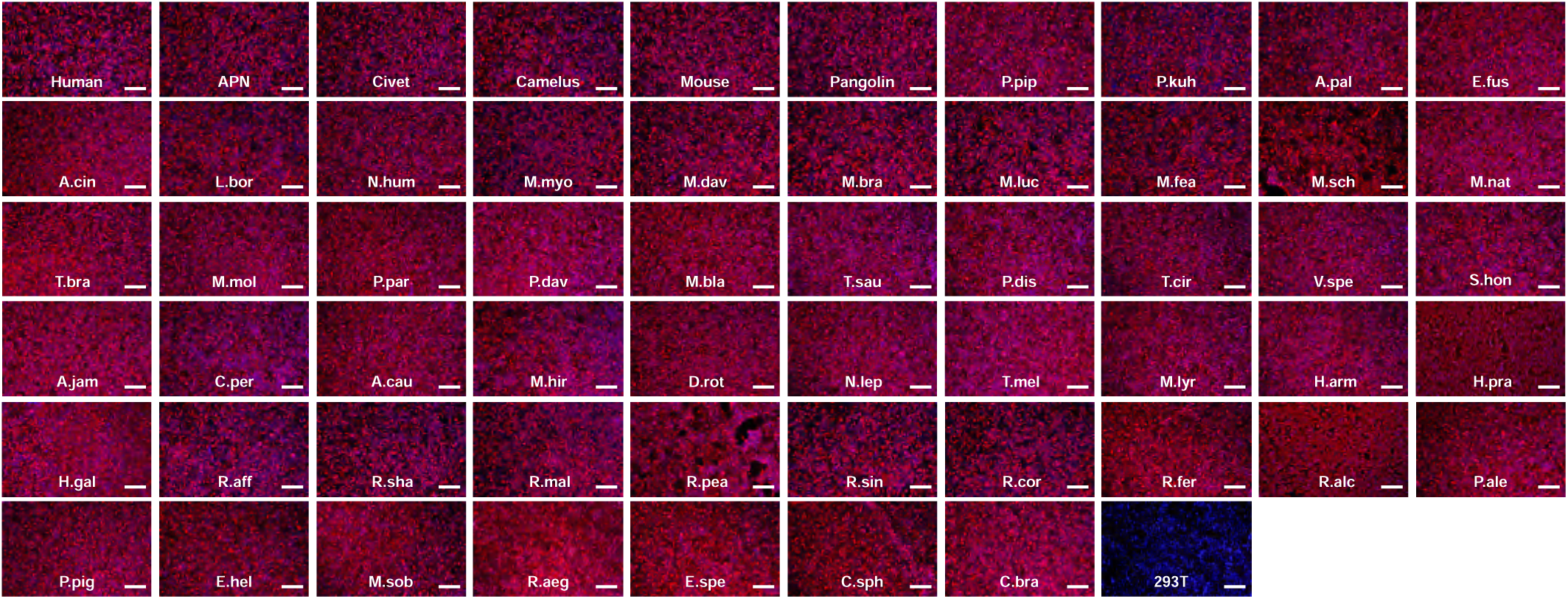
Immunofluorescence demonstrating a comparable expression of 56 ACE2 orthologues in HEK293T cells. The representative images demonstrating the expression of ACE2 orthologues stably expressed in HEK293T cells by detecting the C-terminal fused 3×FLAG tags. The cell nuclei are stained with Hoechst 33342 in blue. Scale bar, 200 μm. Human APN (APN) serves as the experimental control.

**Extended Data Fig. 7.**
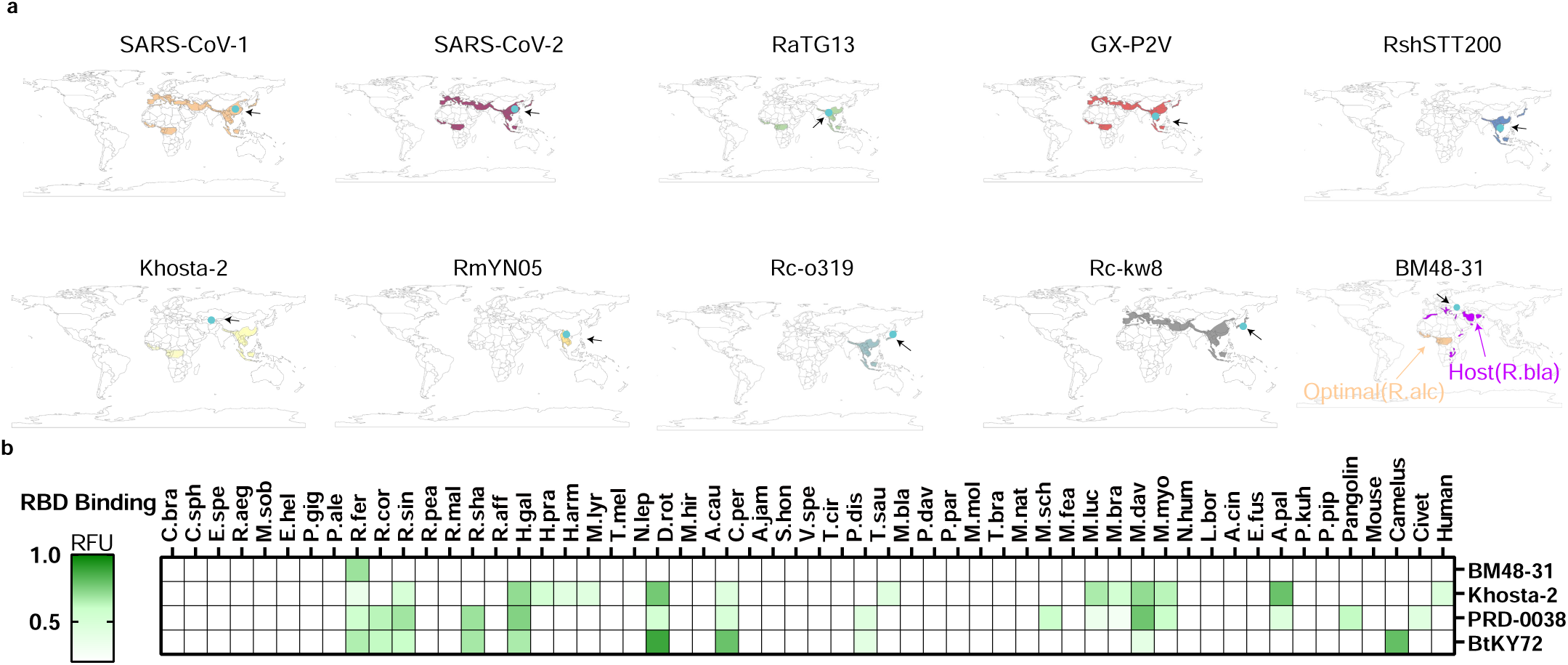
The geographical distribution of Rhinolophus bat species with ACE2 supporting the entry of indicated sarbecovirus (a) and Clade3 sarbecoviruses RBD binding efficiencies with 56 ACE2 orthologues (b). The distribution data are based on ACE2 usage data from Fig. 2b, c. Teal circles: the sampling locations of the sarbecoviruses. For BM48-31, the distribution of the species with the optimal ACE2 orthologue (R.alc) was shown since the sequences of the host (R.bla) ACE2 remains unavailable. RBD-hFc binding was conducted in HEK293T cells expressing the indicated orthologues.

**Extended Data Fig. 8.**
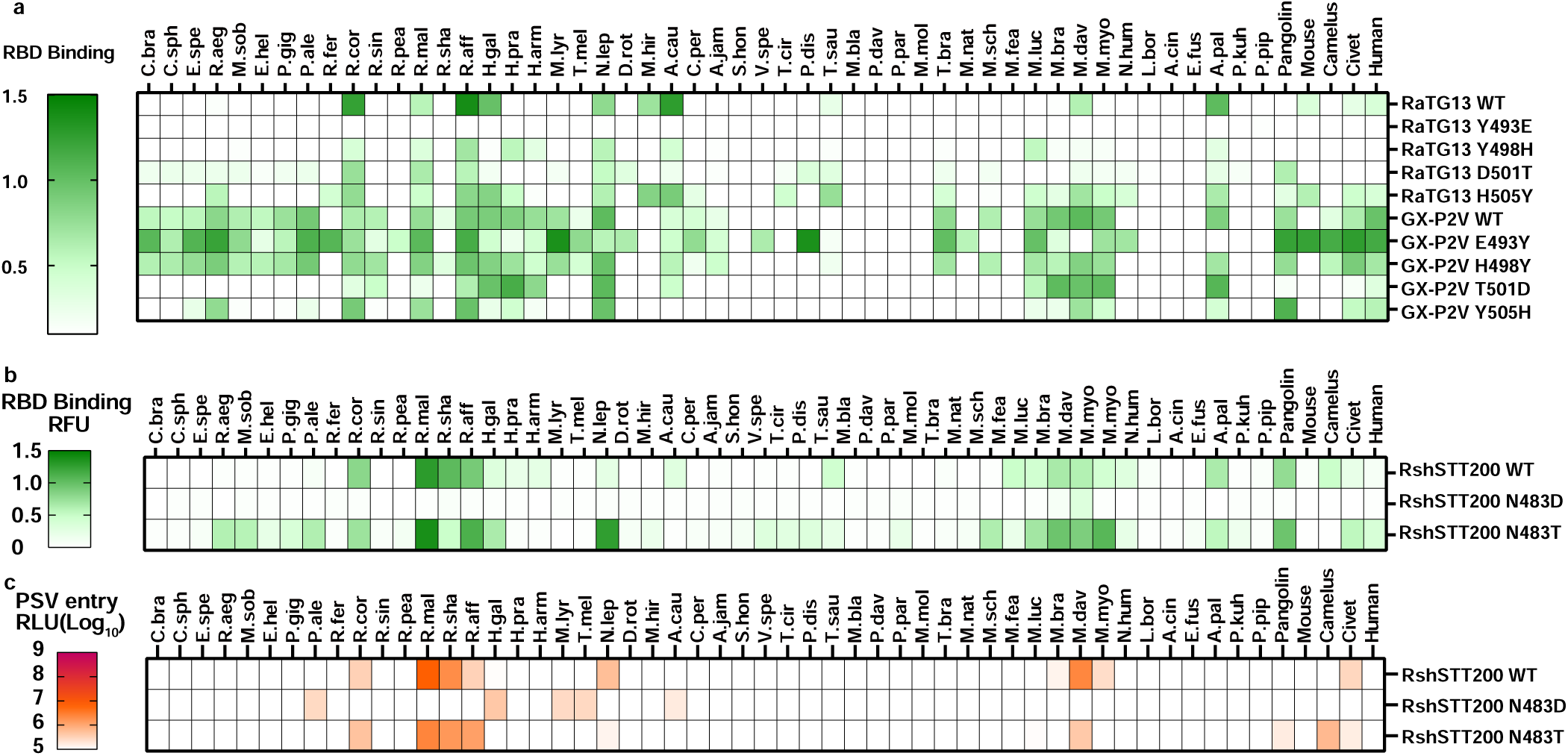
Critical RBM residues affecting the multi-species ACE2 usage spectra of sarbecoviruses. **a**, Heat map displaying the RBD binding efficiencies of RaTG13 and GX-P2V swap mutants at different RBM residues (a) in HEK293T expressing the indicated ACE2 orthologues. **b, c,** Heat map displaying RBD binding(**b**) and PSV entry(**c**) efficiencies of RshSTT200 carrying position 501_SARS-CoV-2_ related mutations.

**Extended Data Fig. 9.**
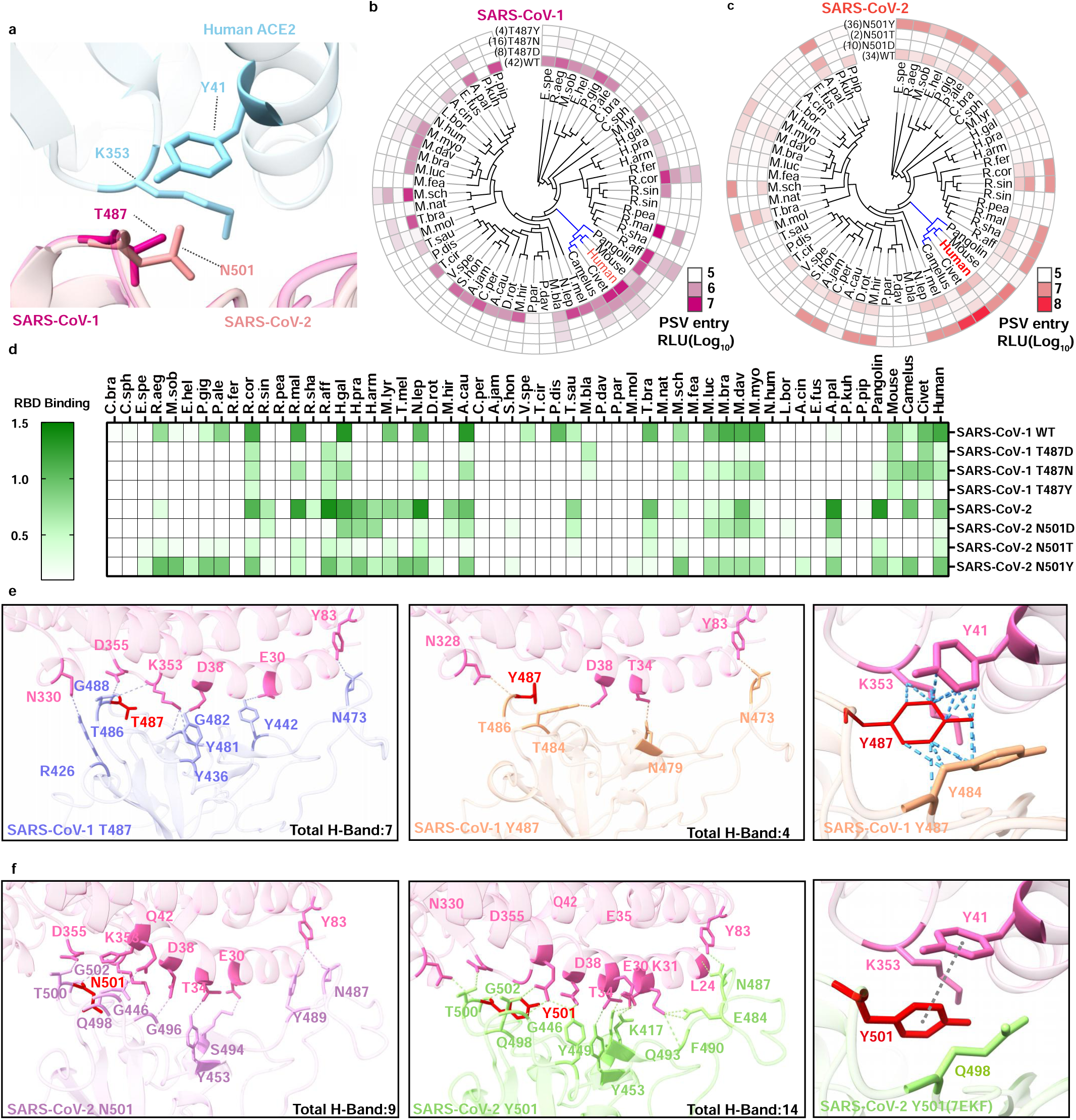
Residue usages in position 501_SARS-CoV-2_ affecting the multi-species ACE2 usage spectra of SARS-CoV-1 and SARS-CoV-2. **a,** Superimposition of the structures illustrating the critical residues for the interaction between SARS-CoV-1/SARS-CoV-2 and human ACE2. **b-d,** Heat map displaying the PSV entry efficiencies of SARS-CoV-1 T487-related mutants (**b**) and SARS-CoV-2 N501-related mutants (**c**), alongside correspond RBD binding (**d)** in HEK293T cells expressing the indicated ACE2 orthologues. PSV entry > 5% is considered as an effective entry and the number of ACE2 support entry is showed in parentheses. **e, f,** Structure modeling of RBD-ACE2 complex illustrating the impact of T487Y_SARS-CoV-1_ (**e**) and N501Y_SARS-CoV-2_ (**f**) mutations on ACE2 interaction, respectively. Blue dashed line: clash. Gray dashed line: π-π stacking interaction.

**Extended Data Fig. 10.**
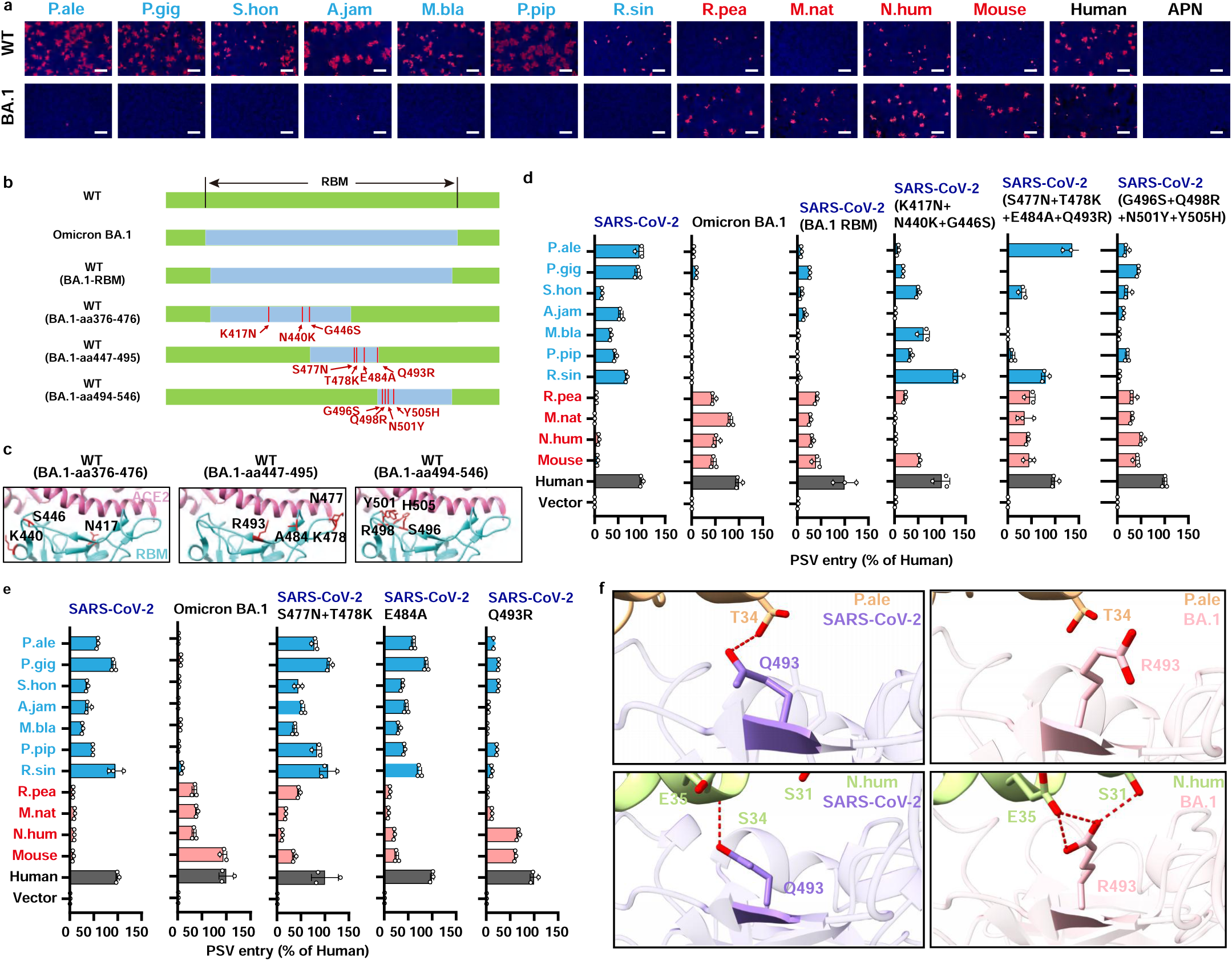
The critical RBM residues responsible for the alteration of multi-species ACE2 usage spectra of Omicron BA.1. **a,** The authentic SARS-CoV-2 (WT) and Omicron BA.1(BA.1) infection in HEK293T stably expressing the ACE2 orthologues showing contrasting entry supporting ability in PSV entry assays (Fig. 2j). Infection efficiencies were examined by immunofluorescence detecting the intracellular N protein. Red font: increased efficiency to support BA.1; Blue font: reduced efficiency to support BA.1. Scale bars, 200 μm. **b,** Schematic diagram illustrating the SARS-CoV-2 mutants with RBM regions swapped between WT and BA.1. **c,** The structural details of the swapped residues within the interaction interface (6M0J). **d, e,** The PSV entry efficiencies of the indicated SARS-CoV-2 mutants carrying region substitutions (**d**) or point mutations (**e**) in HEK293T expressing the indicated ACE2 orthologues. **f,** Structure modeling of SARS-CoV-2 WT or BA.1 RBD in complex with P.ale or N.hum ACE2. The distinct interactions mediated by residue in position 493SARS-CoV-2 were indicated in each model. Red dashed line: H-Bond. The bat ACE2 structures are predicted by Alphafold2 and the complex structures are predicted by HDOCK, SARS-CoV-2 RBD: 6M0J; SARS-CoV-2-BA.1 RBD:7UON.

**Extended Data Fig. 11.**
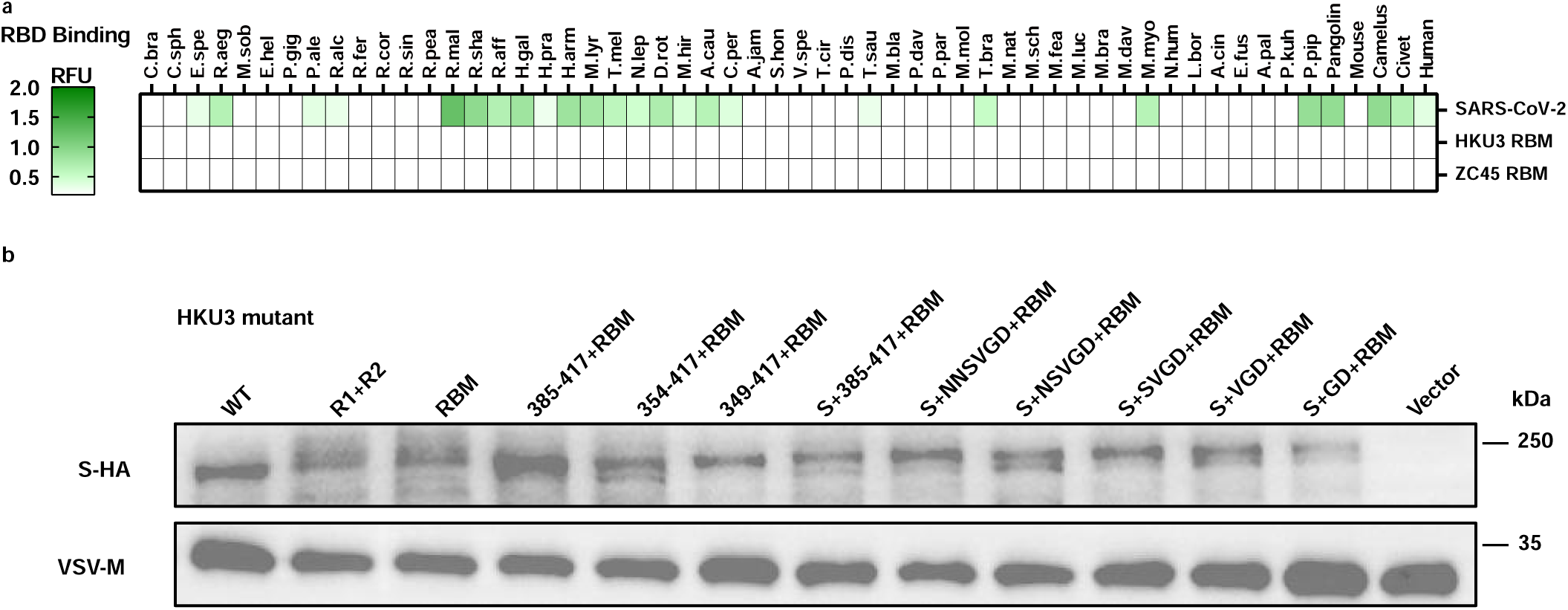
RBD binding efficiency of HKU3 and ZC45 RBM in HEK293T expressing indicated ACE2 orthologues (a) and Western blot detecting the PSV packaging efficiencies of HKU3 (b).

**Extended Data Fig. 12.**
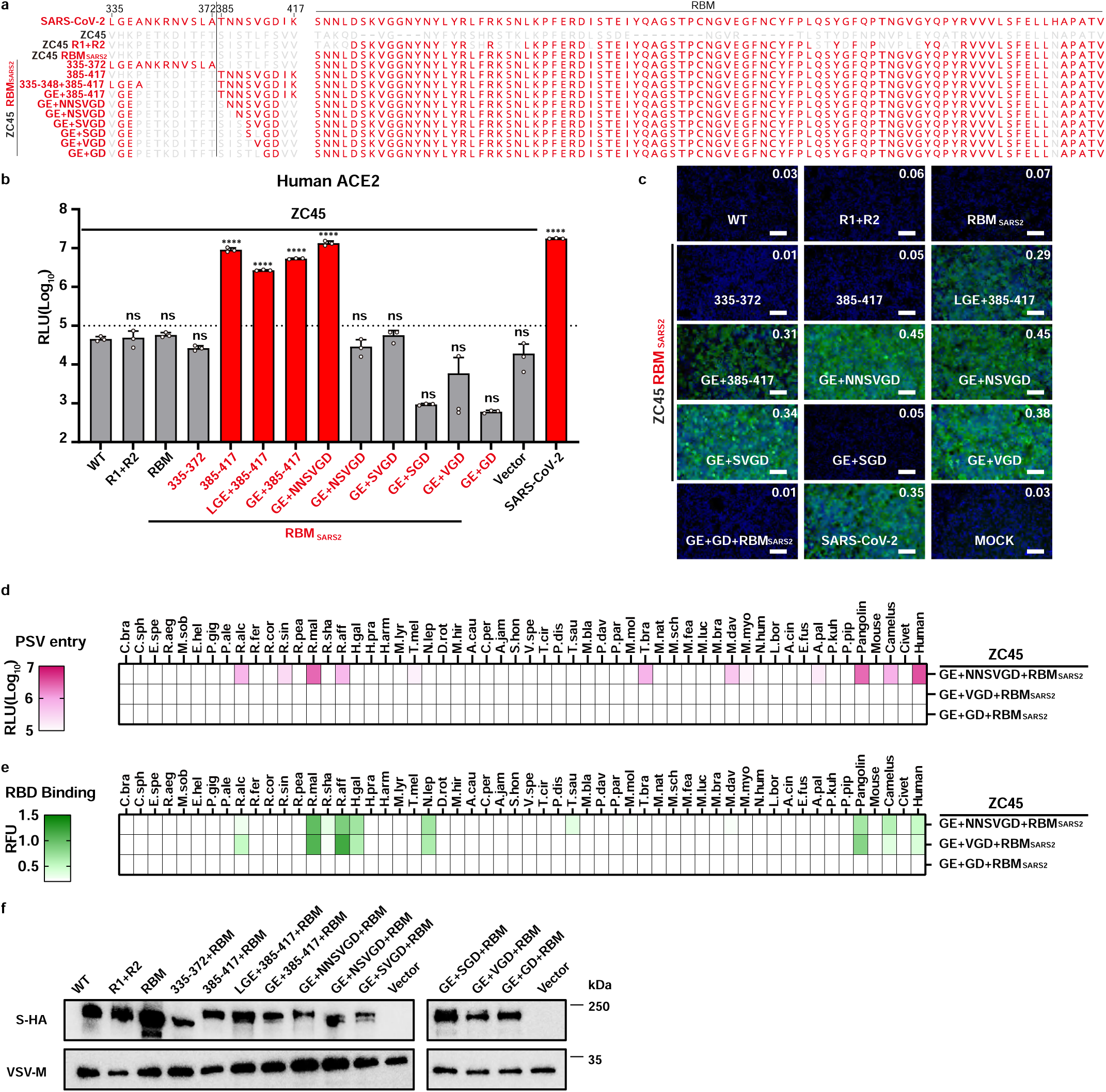
Fine mapping of Clade-2 specific residues outside the ZC45 RBM restricting ACE2 recognition. **a,** Schematic illustration of the mapping strategy to narrow down the critical determinants on ZC45 for ACE2 receptor function. **b, c,** PSV entry (**b**) and RBD binding (**c**) of the ZC45 mutants in HEK293T cells stably expressing hACE2. One-way ANOVA analysis, followed by Dunnett’s test for d and h, mean ± SD. Mock: medium control. Scale bar, 100 μm. GFP RLU is marked on the top right corner. **d, e,**PSV entry (**d**) and RBD binding (**e**) of ZC45 mutants with restored ACE2 binding affinity in HEK293T expressing the indicated ACE2 orthologues. **f**, Western blot illustrating the spike protein package efficiencies of ZC45 mutants in PSV particles.

**Extended Data Fig. 13.**
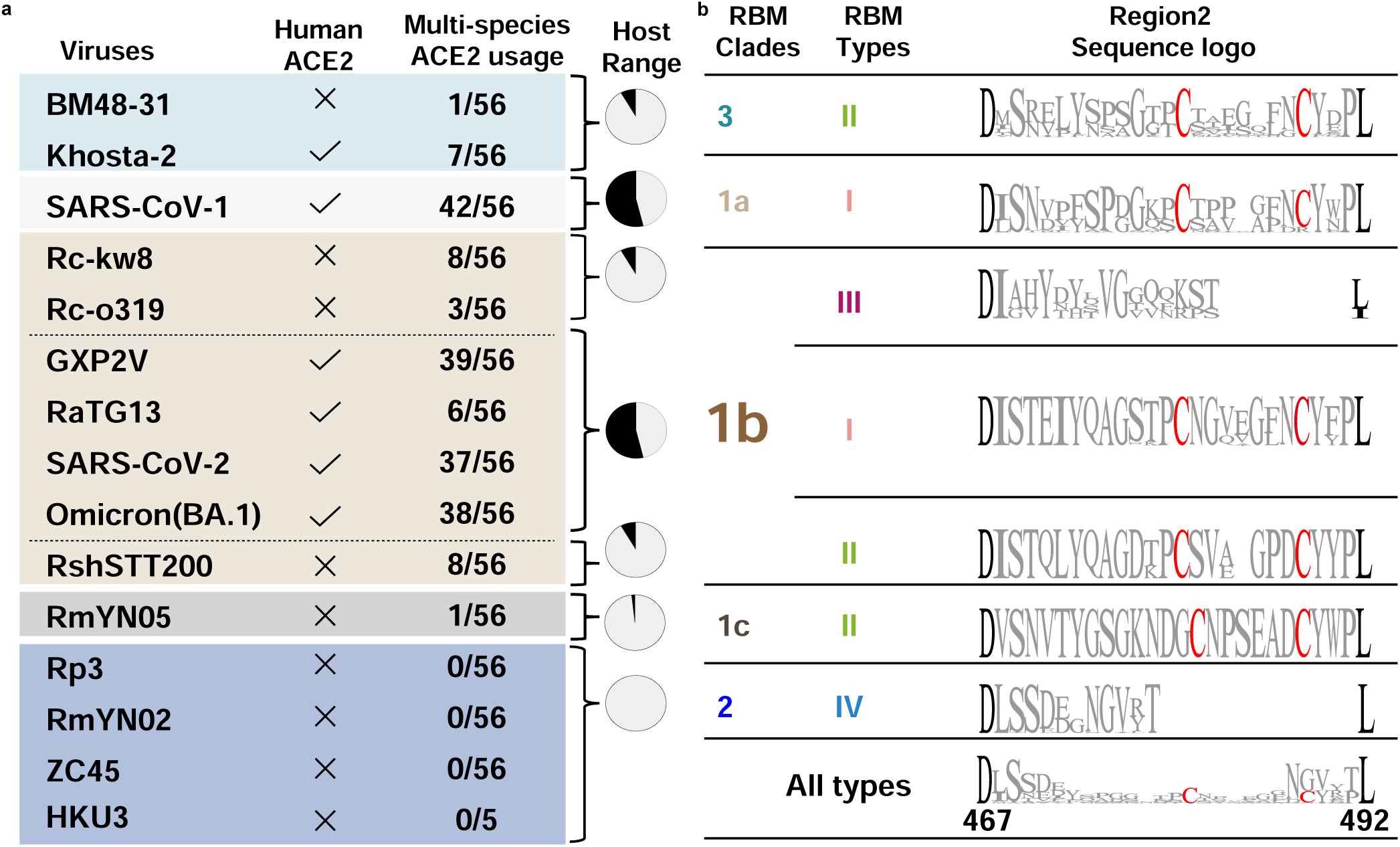
RBM Region 2 sequence logos and ACE2 usage spectra of sarbecoviruses. **a,** Summary of the number of acceptable ACE2 orthologues (data based on Fig.2a and j, RLU>2×10^5^ is considered as positive) and hACE2 compatibility of the indicated sarbecoviruses. Coloring is based on different RBD clades. **b,** Region2 Sequence Logo analysis of sarbecoviruses grouped by different indel types in each clade. The highly conserved D467 and L/I492 (black) (SARS-CoV-2 numbering) for defining the boundary of Region 2 and the featured cystines (red) are highlighted with black and red, respectively.

**Extended Data Fig. 14.**
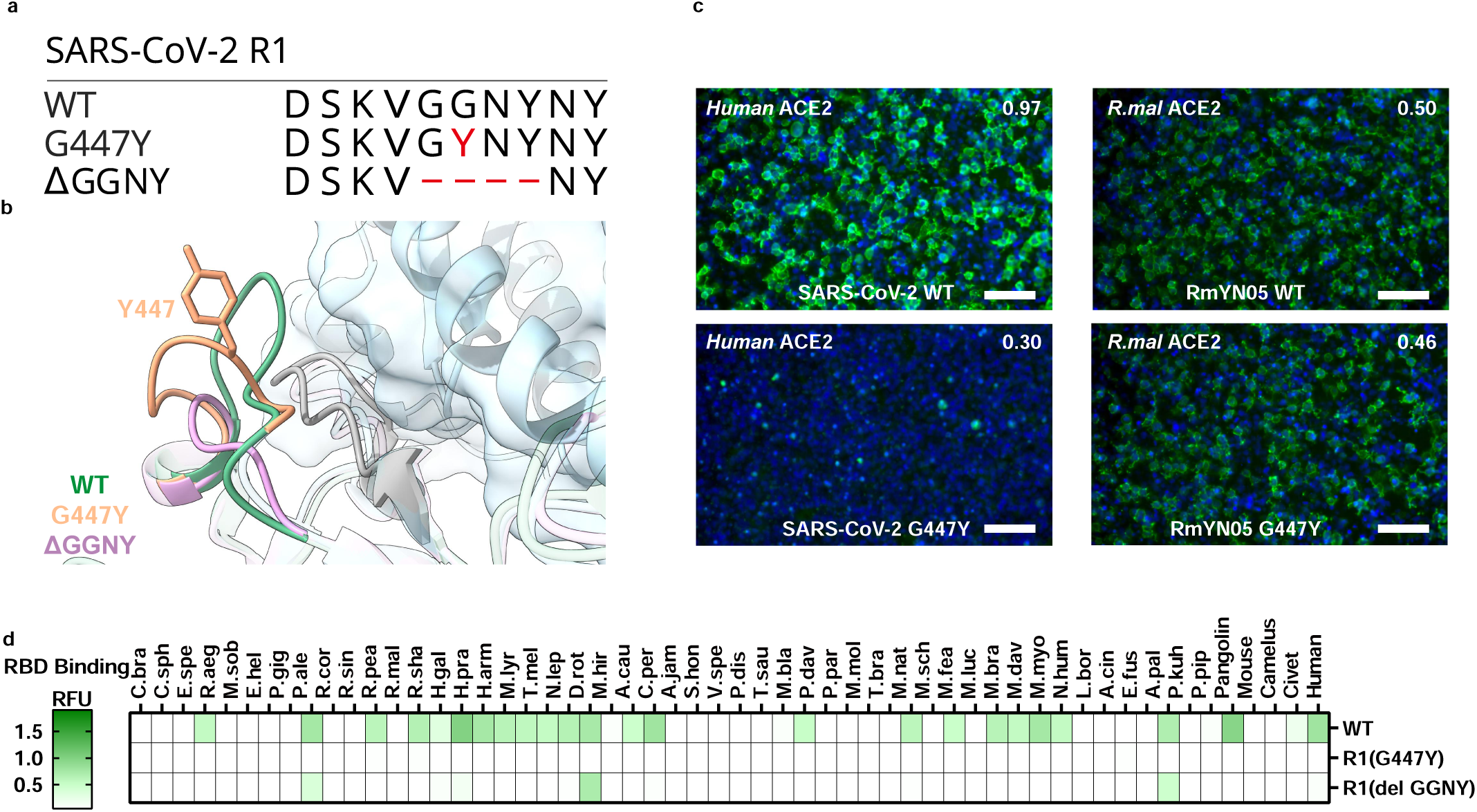
The importance of the conserved RBM Region 1 Glycine (G) on SARS-CoV-2 and ACE2 interaction. **a, b,** Illustration (**a**) and structural modeling (**b**) elucidating the potential impact of SARS-CoV-2 Region 1 mutations on hACE2 interaction (based on 6M0J and AlphaFold2). **c,** The impact of SARS-CoV-2-G447Y mutation and the correspond mutation in RmYN05 on RBD binding efficiencies with host ACE2. Scale bar, 100 μm. **d,** RBD-hFc binding efficiencies of the indicated SARS-CoV-2 RBM Region 1 mutants in HEK293T expressing the indicated ACE2 orthologues.

**Extended Data Fig. 15.**
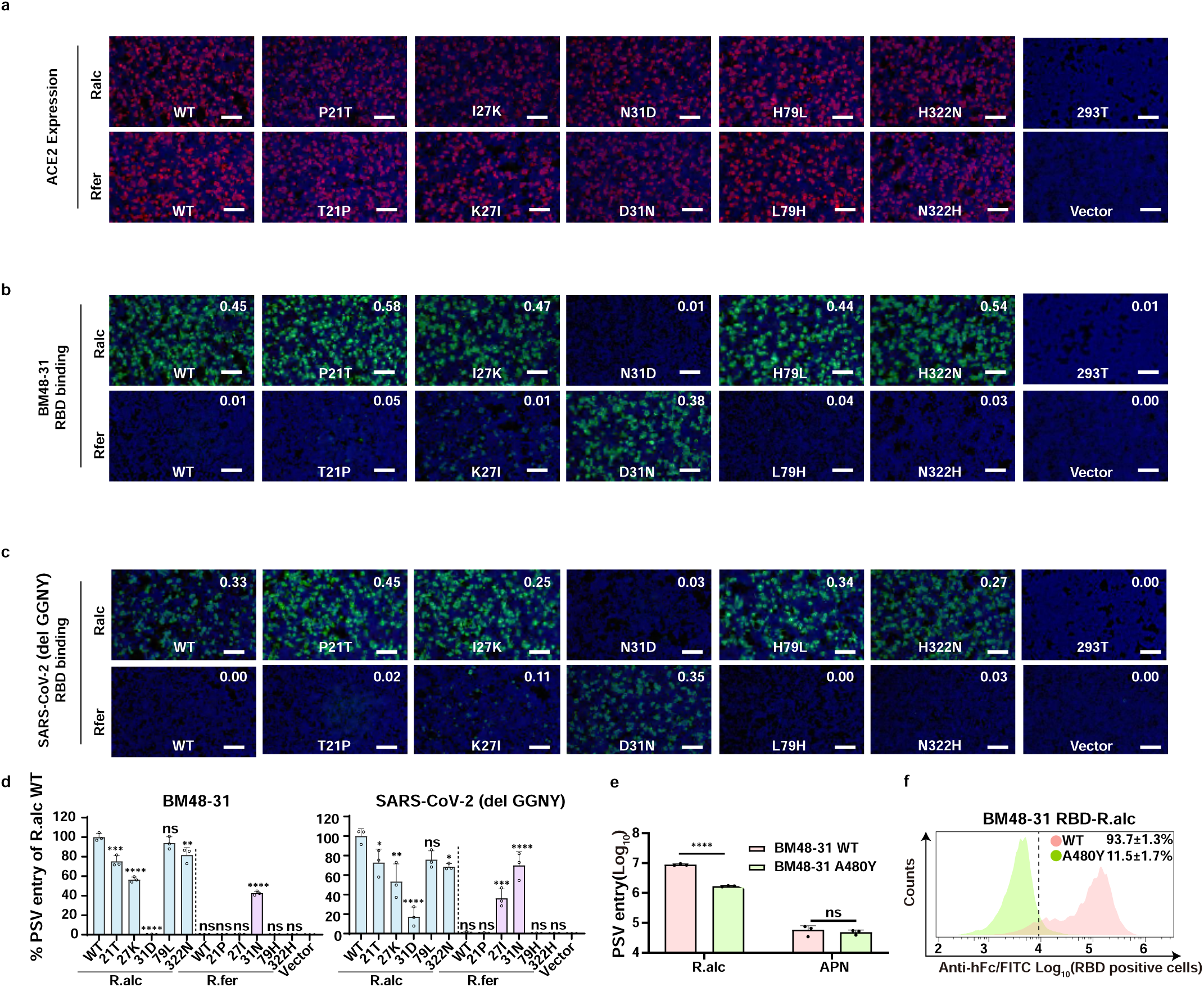
N31_R.alc_ is critical for efficient PSV entry and RBD binding for both BM48-31 and SARS-CoV-2 Δ1(Region1 GGNY deletion). **a,** Expression of R.alc and R.fer ACE2 swap mutants in HEK293T cells by detecting the C-terminal fused FLAG tags. **b-d,** BM48-31(**b**) and SARS-CoV-2 ΔGGNY (**c**) RBD-hFc binding efficiencies and corresponding PSV entry (**d**) in HEK293T expressing indicated R.alc or R.fer ACE2 swap mutants. **e, f,** PSV entry (**e**) and Flow cytometry (**f**) demonstrating the reduced R.alc ACE2 usage upon A480Y_BM48-31_ mutation. *:P<0. 05, **: P<0.01, ***: P<0.001, ****:P<0.0001. The RFU corresponding to each image are indicated on the top right corner. Scale bar: 200 μm. One-way ANOVA analysis, followed by Dunnett’s test, was used for statistical analysis of significance. Two-tailed unpaired (Student’s) t-test was performed if only two conditions were compared. Bar charts presented in mean ± s.d.

## Graphic Abstract

**Extended Data Fig. 16.**
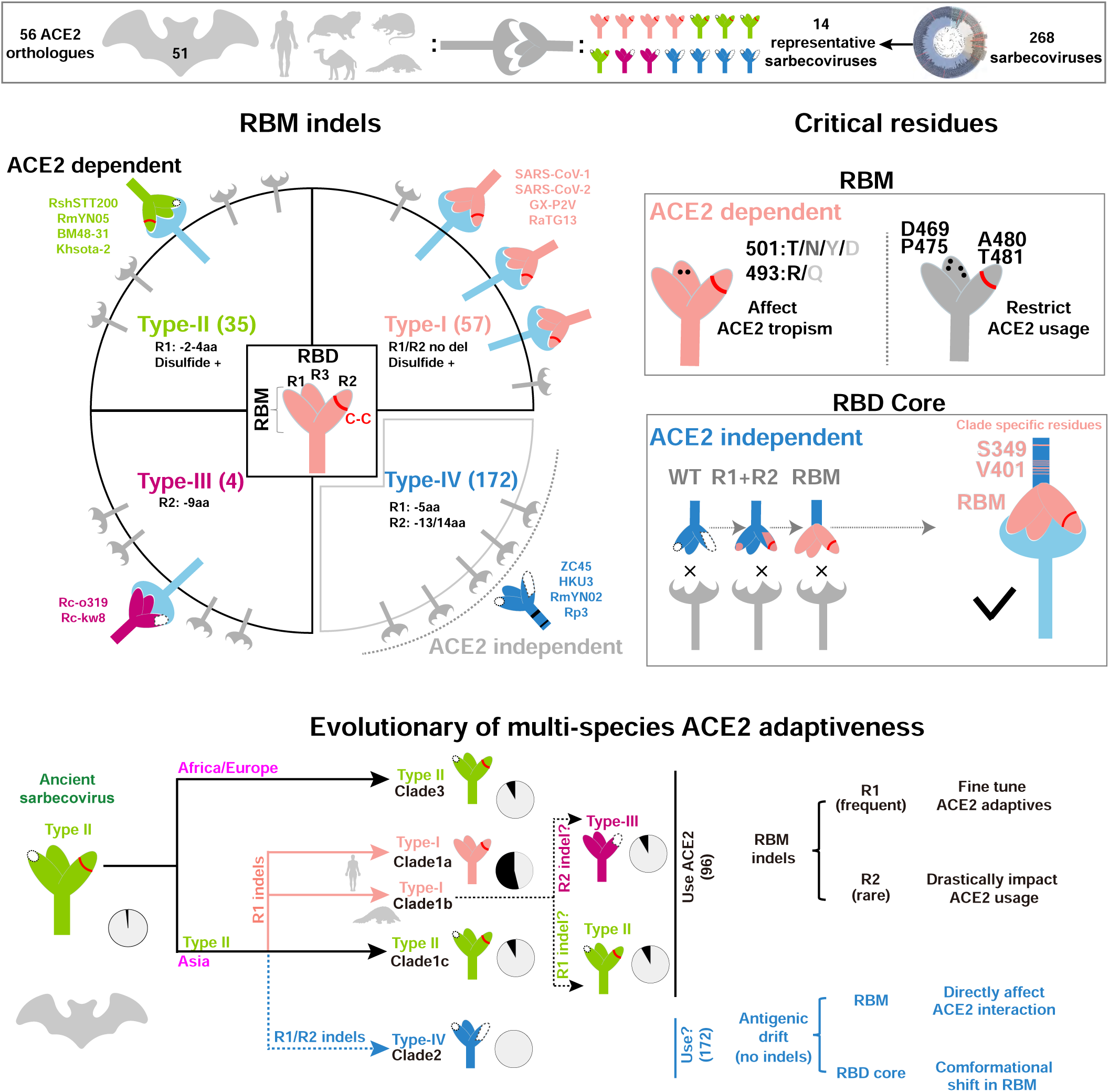
Graphic Abstract. Multi-species ACE2 tropism of sarbecoviruses of different RBM types and the proposed evolutionary of their multi-species ACE2 adaptiveness.

## References

1. Starr, T. N. et al. ACE2 binding is an ancestral and evolvable trait of sarbecoviruses. Nature 603, 913–918 (2022).

2. Hu, B. et al. Discovery of a rich gene pool of bat SARS-related coronaviruses provides new insights into the origin of SARS coronavirus. PLoS Pathog. 13, e1006698 (2017).

3. Letko, M., Marzi, A. & Munster, V. Functional assessment of cell entry and receptor usage for SARS-CoV-2 and other lineage B betacoronaviruses. Nat. Microbiol. 5, 562–569 (2020).

4. Murakami, S. et al. Isolation of Bat Sarbecoviruses, Japan. Emerg. Infect. Dis. 28, 2500–2503 (2022).

5. Guo, H. et al. Identification of a novel lineage bat SARS-related coronaviruses that use bat ACE2 receptor. Emerg. Microbes Infect. 10, 1507–1514 (2021).

6. Zhou, H. et al. Identification of novel bat coronaviruses sheds light on the evolutionary origins of SARS-CoV-2 and related viruses. Cell 184, 4380–4391.e14 (2021).

7. Wacharapluesadee, S. et al. Evidence for SARS-CoV-2 related coronaviruses circulating in bats and pangolins in Southeast Asia. Nat. Commun. 12, 972 (2021).

8. Alkhovsky, S. et al. SARS-like Coronaviruses in Horseshoe Bats (Rhinolophus spp.) in Russia, 2020. Viruses 14, 113 (2022).

9. Lam, T. T.-Y. et al. Identifying SARS-CoV-2-related coronaviruses in Malayan pangolins. Nature 583, 282–285 (2020).

10. Delaune, D. et al. A novel SARS-CoV-2 related coronavirus in bats from Cambodia. Nat. Commun. 12, 6563 (2021).

11. Guo, H. et al. ACE2-Independent Bat Sarbecovirus Entry and Replication in Human and Bat Cells. mBio 13, e0256622 (2022).

12. Ksiazek, T. G. et al. A novel coronavirus associated with severe acute respiratory syndrome. N. Engl. J. Med. 348, 1953–1966 (2003).

13. Zhou, P. et al. A pneumonia outbreak associated with a new coronavirus of probable bat origin. Nature 579, 270–273 (2020).

14. Meta Djomsi, D., et al. Coronaviruses Are Abundant and Genetically Diverse in West and Central African Bats, including Viruses Closely Related to Human Coronaviruses. Viruses 15, 337 (2023).

15. Murakami, S. et al. Detection and Characterization of Bat Sarbecovirus Phylogenetically Related to SARS-CoV-2, Japan. Emerg. Infect. Dis. 26, 3025–3029 (2020).

16. Temmam, S. et al. Bat coronaviruses related to SARS-CoV-2 and infectious for human cells. Nature 604, 330– 336 (2022).

17. Ge, X.-Y. et al. Isolation and characterization of a bat SARS-like coronavirus that uses the ACE2 receptor. Nature 503, 535–538 (2013).

18. Li, W. et al. Bats are natural reservoirs of SARS-like coronaviruses. Science 310, 676–679 (2005).

19. Wu, Z. et al. A comprehensive survey of bat sarbecoviruses across China in relation to the origins of SARS-CoV and SARS-CoV-2. Natl. Sci. Rev. 10, nwac213 (2023).

20. Xiao, K. et al. Isolation of SARS-CoV-2-related coronavirus from Malayan pangolins. Nature 583, 286–289 (2020).

21. Guo, Z. et al. SARS-CoV-2-related pangolin coronavirus exhibits similar infection characteristics to SARS-CoV-2 and direct contact transmissibility in hamsters. iScience 25, 104350 (2022).

22. Niu, S. et al. Molecular basis of cross-species ACE2 interactions with SARS-CoV-2-like viruses of pangolin origin. EMBO J. 40, (2021).

23. Li, P. et al. The Rhinolophus affinis bat ACE2 and multiple animal orthologs are functional receptors for bat coronavirus RaTG13 and SARS-CoV-2. Sci. Bull. 66, 1215–1227 (2021).

24. Liu, K. et al. Binding and molecular basis of the bat coronavirus RaTG13 virus to ACE2 in humans and other species. Cell 184, 3438–3451.e10 (2021).

25. Boni, M. F. et al. Evolutionary origins of the SARS-CoV-2 sarbecovirus lineage responsible for the COVID-19 pandemic. Nat. Microbiol. 5, 1408–1417 (2020).

26. Hofmann, H. et al. Human coronavirus NL63 employs the severe acute respiratory syndrome coronavirus receptor for cellular entry. Proc. Natl. Acad. Sci. 102, 7988–7993 (2005).

27. Xiong, Q. et al. Close relatives of MERS-CoV in bats use ACE2 as their functional receptors. Nature 612, 748– 757 (2022).

28. Lan, J. et al. Structure of the SARS-CoV-2 spike receptor-binding domain bound to the ACE2 receptor. Nature 581, 215–220 (2020).

29. Wu, K., Peng, G., Wilken, M., Geraghty, R. J. & Li, F. Mechanisms of Host Receptor Adaptation by Severe Acute Respiratory Syndrome Coronavirus. J. Biol. Chem. 287, 8904 (2012).

30. Yan, H. et al. ACE2 receptor usage reveals variation in susceptibility to SARS-CoV and SARS-CoV-2 infection among bat species. *Nat*. Ecol. Evol. 5, 600–608 (2021).

31. Liu, Y., et al. Functional and genetic analysis of viral receptor ACE2 orthologs reveals a broad potential host range of SARS-CoV-2. Proc. Natl. Acad. Sci. 118, e2025373118 (2021).

32. Zhang, Y. et al. Cross-species tropism and antigenic landscapes of circulating SARS-CoV-2 variants. Cell Rep. 38, 110558 (2022).

33. Li, P. et al. Effect of polymorphism in Rhinolophus affinis ACE2 on entry of SARS-CoV-2 related bat coronaviruses. PLOS Pathog. 19, e1011116 (2023).

34. Guo, H. et al. Evolutionary Arms Race between Virus and Host Drives Genetic Diversity in Bat Severe Acute Respiratory Syndrome-Related Coronavirus Spike Genes. J. Virol. 94, e00902–20 (2020).

35. Wells, H. L. et al. The evolutionary history of ACE2 usage within the coronavirus subgenus *Sarbecovirus*. Virus Evol. 7, veab007 (2021).

36. Roelle, S. M., Shukla, N., Pham, A. T., Bruchez, A. M. & Matreyek, K. A. Expanded ACE2 dependencies of diverse SARS-like coronavirus receptor binding domains. PLOS Biol. 20, e3001738 (2022).

37. Hu, D. et al. Genomic characterization and infectivity of a novel SARS-like coronavirus in Chinese bats. Emerg. Microbes Infect. 7, 154 (2018).

38. Ren, W. et al. Difference in receptor usage between severe acute respiratory syndrome (SARS) coronavirus and SARS-like coronavirus of bat origin. J. Virol. 82, 1899–1907 (2008).

39. Crook, J. M. et al. Metagenomic identification of a new sarbecovirus from horseshoe bats in Europe. Sci. Rep. 11, 14723 (2021).

40. Drexler, J. F. et al. Genomic characterization of severe acute respiratory syndrome-related coronavirus in European bats and classification of coronaviruses based on partial RNA-dependent RNA polymerase gene sequences. J. Virol. 84, 11336–11349 (2010).

41. Tao, Y. & Tong, S. Complete Genome Sequence of a Severe Acute Respiratory Syndrome-Related Coronavirus from Kenyan Bats. Microbiol. Resour. Announc. 8, e00548–19 (2019).

42. Lee, J. et al. Broad receptor tropism and immunogenicity of a clade 3 sarbecovirus. Cell Host Microbe 31, 1961–1973.e11 (2023).

43. Seifert, S. N. et al. An ACE2-dependent Sarbecovirus in Russian bats is resistant to SARS-CoV-2 vaccines. PLOS Pathog. 18, e1010828 (2022).

44. Zech, F. et al. Spike residue 403 affects binding of coronavirus spikes to human ACE2. Nat. Commun. 12, 6855 (2021).

45. Grishin, A. M. et al. Disulfide Bonds Play a Critical Role in the Structure and Function of the Receptor-binding Domain of the SARS-CoV-2 Spike Antigen. J. Mol. Biol. 434, 167357 (2022).

46. Wang, Q. et al. Key mutations on spike protein altering ACE2 receptor utilization and potentially expanding host range of emerging SARS-CoV-2 variants. J. Med. Virol. 95, e28116 (2023).

47. Zhao, Z. et al. Structural basis for receptor binding and broader interspecies receptor recognition of currently circulating Omicron sub-variants. Nat. Commun. 14, 4405 (2023).

48. Starr, T. N. et al. Shifting mutational constraints in the SARS-CoV-2 receptor-binding domain during viral evolution. Science 377, 420–424 (2022).

49. Han, P. et al. Molecular insights into receptor binding of recent emerging SARS-CoV-2 variants. Nat. Commun. 12, 6103 (2021).

50. Hati, S. & Bhattacharyya, S. Impact of Thiol–Disulfide Balance on the Binding of Covid-19 Spike Protein with Angiotensin-Converting Enzyme 2 Receptor. ACS Omega 5, 16292–16298 (2020).

51. Gao, B. & Zhu, S. Mutation-driven parallel evolution in emergence of ACE2-utilizing sarbecoviruses. Front. Microbiol. 14, 1118025 (2023).

52. Mykytyn, A. Z., Fouchier, R. A. & Haagmans, B. L. Antigenic evolution of SARS coronavirus 2. Curr. Opin. Virol. 62, 101349 (2023).

53. Menachery, V. D. et al. Trypsin Treatment Unlocks Barrier for Zoonotic Bat Coronavirus Infection. J. Virol. 94, e01774–19 (2020).

54. Richard, M. et al. Factors determining human-to-human transmissibility of zoonotic pathogens via contact. Curr. Opin. Virol. 22, 7–12 (2017).

## References

55. Nakamura, T., Yamada, K. D., Tomii, K. & Katoh, K. Parallelization of MAFFT for large-scale multiple sequence alignments. Bioinforma. Oxf. Engl. 34, 2490–2492 (2018).

56. Nguyen, L.-T., Schmidt, H. A., von Haeseler, A. & Minh, B. Q. IQ-TREE: a fast and effective stochastic algorithm for estimating maximum-likelihood phylogenies. Mol. Biol. Evol. 32, 268–274 (2015).

57. Schwegmann-Weßels, C. et al. Comparison of vesicular stomatitis virus pseudotyped with the S proteins from a porcine and a human coronavirus. J. Gen. Virol. 90, 1724–1729 (2009).

58. Fukushi, S. et al. Vesicular stomatitis virus pseudotyped with severe acute respiratory syndrome coronavirus spike protein. J. Gen. Virol. 86, 2269–2274 (2005).

59. Ma, C. et al. Broad host tropism of ACE2-using MERS-related coronaviruses and determinants restricting viral recognition. Cell Discov. 9, 57 (2023).

60. Nie, J. et al. Quantification of SARS-CoV-2 neutralizing antibody by a pseudotyped virus-based assay. Nat. Protoc. 15, 3699–3715 (2020).

61. Whitt, M. A. Generation of VSV pseudotypes using recombinant ΔG-VSV for studies on virus entry, identification of entry inhibitors, and immune responses to vaccines. J. Virol. Methods 169, 365–374 (2010).

62. Jumper, J. et al. Highly accurate protein structure prediction with AlphaFold. Nature 596, 583–589 (2021).

63. Yan, Y., Tao, H., He, J. & Huang, S.-Y. The HDOCK server for integrated protein-protein docking. Nat. Protoc. 15, 1829–1852 (2020).

64. Yan, Y., Wen, Z., Wang, X. & Huang, S.-Y. Addressing recent docking challenges: A hybrid strategy to integrate template-based and free protein-protein docking. Proteins 85, 497–512 (2017).

