## Supplementary figures and images for "Sarbecovirus RBD indels and specific residues dictating ACE2 multi-species adaptiveness"

### Supplementary data S2

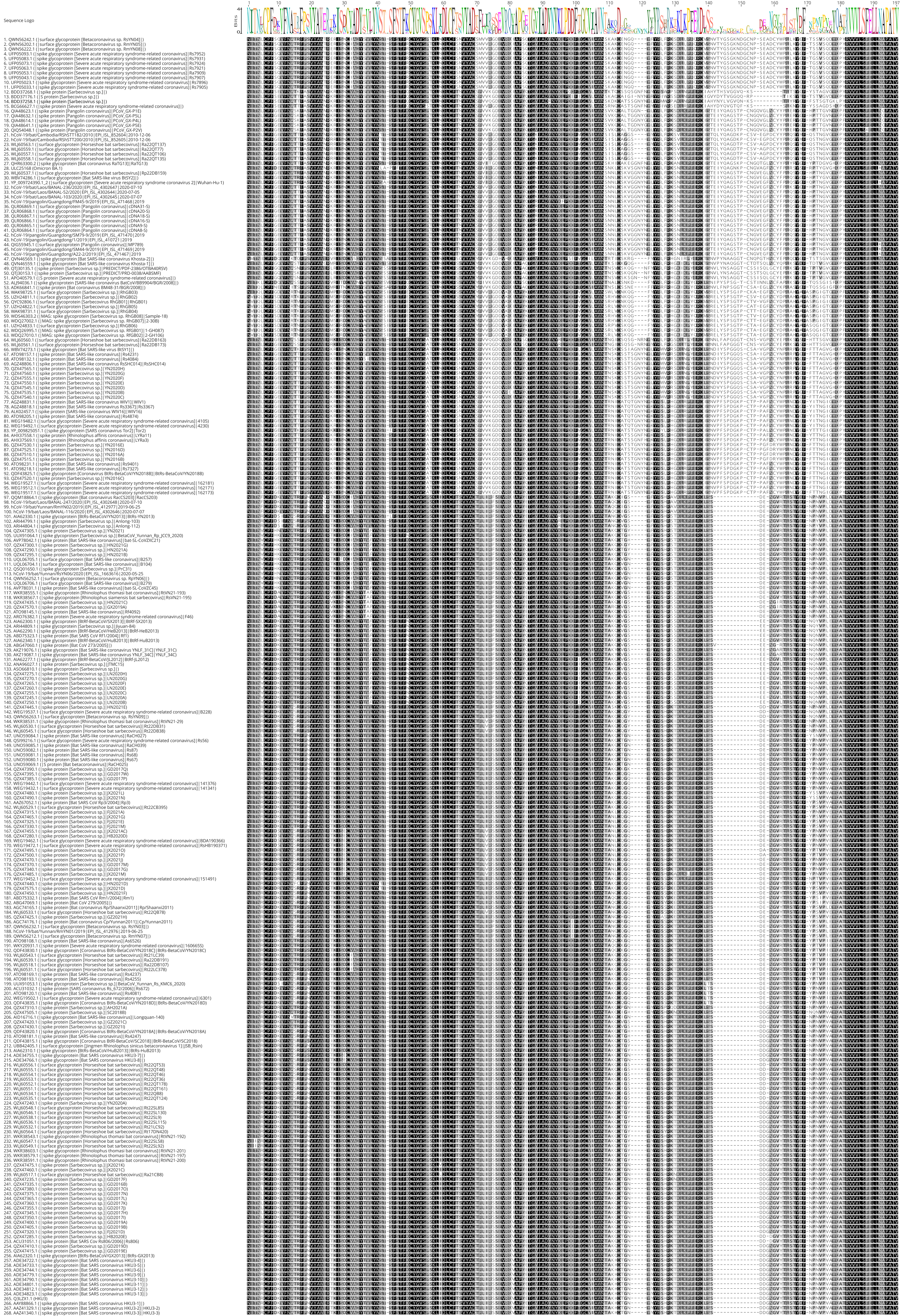
